# Specificity of the stabilising interaction between intrinsically disordered protein sequences and G-quadruplexes in RNA

**DOI:** 10.1101/2025.10.19.683336

**Authors:** Lachlan B. Cox, John S. Mattick, Isis A. Middleton, Felix J. Rizzuto, Pall Thordarson

## Abstract

Intrinsically disordered regions (IDRs) are present in and essential for the function of nearly all the proteins involved in the regulation, cell and developmental processes. The RGG domain in IDRs binds ‘promiscuously’ to RNA G-quadruplexes (rG4s), a non-canonical 4-stranded secondary structure that occurs in many transcripts involved in gene regulation. Here we show, using weak binding interactions between a minimal RGG-rich peptide and rG4s, that the IDR selectively templates and stabilises the structure of the human telomeric TERRA rG4, providing a unique pathway to RNA folding that does not rely on high-affinity binding or monovalent cations. Multidimensional NMR and circular dichroism analyses reveal individual nucleotide and amino acid identities determine the specificity of the interaction between RGG peptides and rG4s, explaining how IDRs can selectively recognise RNA over DNA G4s, the high specificity of such interactions *in vivo,* and the high frequency of monogenic mutations observed in IDRs.

**GRAPHICAL ABSTRACT:** 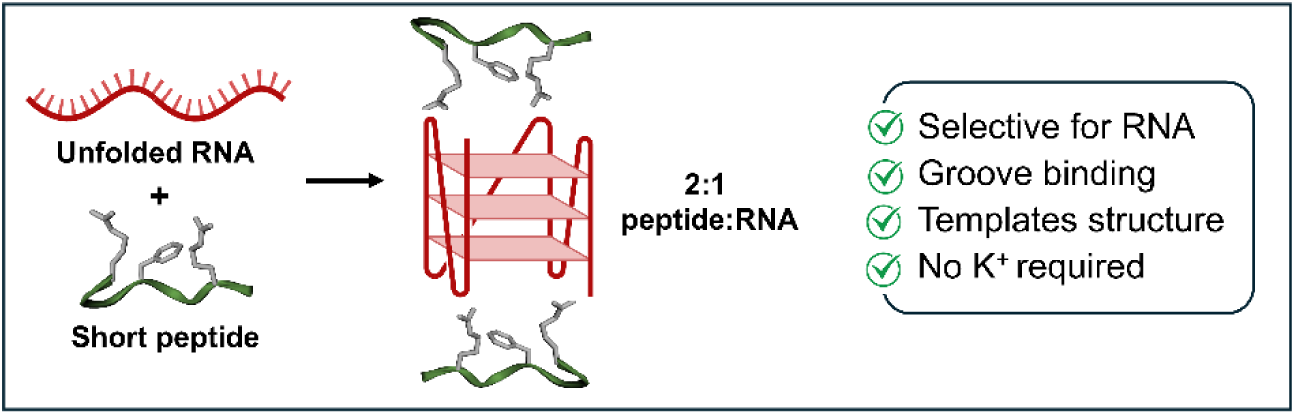

## INTRODUCTION

Nucleic acids form diverse secondary structures, many of which modulate cellular functions (1). Apart from the canonical Watson-Crick duplex, DNA and RNA can form non-canonical structures under specific conditions (2–4). Among these, G-quadruplexes (G4s) are the most prevalent folded non-canonical structures in the genome (5). These motifs are thought to be involved in genetic disorders by altering transcriptional, translational and epigenetic regulatory factors (6), resulting in neurodegenerative diseases (7), myotonic dystrophy, and cancers (8). G4s are formed of planar guanine tetrads, held together by self-complementary hydrogen bonding templated around a central monovalent cation (Figure 1C) (9), and occur in a wide variety of mRNAs and long noncoding RNAs (lncRNAs), including those involved telomere maintenance, enhancer action, modulation of chromatin structure, transcription, RNA splicing, RNA transport and translation (10–12). The termini of human chromosomes – the telomere – contains instructions for this structure in cells, with capping regions of 300-8000 tandem repeats of 5’-CCCTAA/TTAGGG-3’ (13). This region is tasked with regulating telomere integrity upon interaction with proteins (14). When transcribed, the telomeric repeat-containing, non-coding RNA (TERRA) sequence (GGG UUA)_n_ is formed, folding into a parallel G4 structure (15).

**Figure 1.**
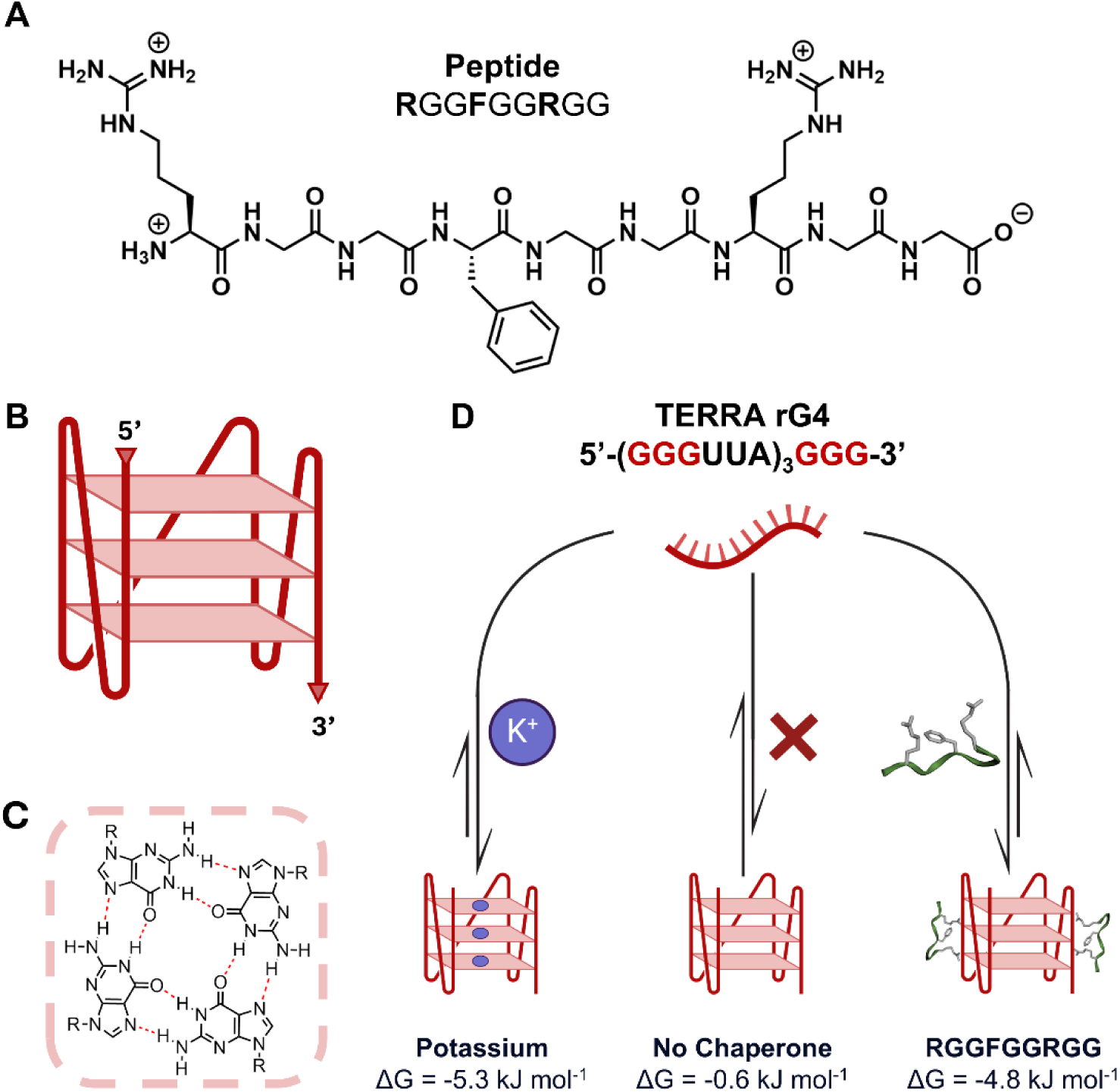
Peptide and oligonucleotide sequences used in this study. **(A)**: Sequence of the RNA-binding peptide **RGGFGGRGG**, derived from the nucleolin intrinsically disordered region. **(B)**: Folded structure of the TERRA rG4 structure. **(C)**: Four guanines can interact via Hoogsteen hydrogen bonding to form a tetrad sheet. (**D)**: The rG4 structure can be chaperoned by either potassium ions (left), or the **RGGFGGRGG** peptide (right) to a comparable magnitude. The free energy of templation (ΔG) at 25 °C is shown below each structure. The 5’ to 3’ RNA phosphate backbone is represented by thick lines and the Hoogsteen base pairs are indicated by thin horizontal sheets.

G4s interact strongly with the intrinsically disordered regions (IDRs) of proteins (16), long chains of amino acids that lack rigid tertiary structure and are characterised by a high proportion of small, polar and positively charged amino acids (glycine, arginine, histidine and lysine), often in the form of RGG/RG, histidine-rich domains or other repeats (17–20). IDRs are essential for the function of nearly all of the proteins involved in gene regulation, including RNA polymerase, most transcription factors, enhancers, Hox proteins, histones, histone modifying proteins, other chromatin-binding proteins, the Mediator complex, RNA binding proteins, splicing factors, membrane receptors, cytoskeletal proteins and nuclear hormone receptors (21–35). IDRs are also major sites of post-translational modifications and interact with a multitude of RNAs in the formation of biomolecular condensates or ‘phase-separated domains’, in the nucleus and cytoplasm (36–38).

Like G4 quadruplexes, IDRs are described as ‘promiscuous’, i.e., they interact with many partners (19,20,26,39–51). The aberrant promiscuity of IDR-containing proteins and perturbations of phase-separated domains may underlie the dosage sensitivity of oncogenes and other proteins (52,53) as well as neurodegenerative disorders (54).

One of the most common RNA-binding motifs in IDRs is the RGG domain, which binds to G-quadruplexes in RNAs (rG4s), reciprocally mediating degenerate specificity (55). The specificity of the RGG-rG4 interaction is poorly understood. A previous study established that multiple hydrogen bonds are involved in the interaction of an RGG peptide from the human fragile X mental retardation protein (FMRP) and an *in vitro*–selected guanine-rich RNA, validated by RNA and peptide mutations (56). The IDR-directed binding of two different transcription factors *in vivo* was also shown to be due to weak determinants distributed throughout the IDR sequence (47) and that phenylalanines adjacent to RGG motifs in the IDR of nucleolin are responsible for rG4 binding and folding (57). Despite the importance of IDR-RNA interactions in biological systems, however, no studies have thus far confirmed the role of specific intermolecular interactions in stabilizing, forming or disrupting non-canonical RNA secondary structures with proteins rich in IDRs (12,57,58). To gain a holistic understanding of the nature and specificity of RNA-protein interactions in the formation of biocondensates,(59) and the perturbation of these interactions in disease, unravelling the relatively weak interactions contributing to IDR-RNA binding is key.

Here we analyse in detail the interactions between the peptide **RGGFGGRGG**, derived from the IDR of the nucleolin RNA binding protein (Figure 1A), and folded G4 structures from the human telomeric repeat-containing lncRNA (TERRA) sequence [5’-(GGG UUA GGG UUA GGG UUA GGG-3’] (Figure 1B) (15,60). We show that this peptide stabilises short natural sequences of human telomeric RNA to form an rG4 secondary structure, despite a relatively modest binding affinity. The peptide discriminates binding RNAs at a single-nucleotide mutation, is selective for RNA over DNA, and is competitive with the endogenous chaperone K^+^ (Figure 1D). Mutation of the peptide sequence abrogates or severely weakens its binding, highlighting the essential amino acid residues of arginine and phenylalanine required for RNA stabilisation. Our results are consistent with the observation that mutations that cause human monogenic diseases occur more commonly in IDRs than globular domains (61,62), indicating that, despite their relatively simple composition, IDRs have strong sequence constraints.

## MATERIALS AND METHODS

### Solid-phase peptide synthesis and purification

The peptides used in this study were synthesised through standard solid-phase peptide synthesis (Supplementary Information Section 2.2). The purification was performed on a Shimadzu Prominence UFLC HPLC fitted with a Shimadzu 20AB pump and an SPD-20A photodiode array detector. An Intersil ODS-4 5 μm 20 x 150 mm column with a C18 stationary phase was used with mobile phases A: MilliQ™ water with formic acid (0.1%, v/v), and B: acetonitrile with formic acid (0.1%, v/v). The concentration of mobile phase B increased from 10-95% over 35 min, with a flow rate of 5 mL min^-1^ and monitored at 254 nm. Samples were lyophilised to afford purified peptide as a white powder.

### Solid phase oligonucleotide synthesis and purification

RNA oligonucleotides were synthesised by the UNSW RNA Institute on an ÄKTA Oligopilot Plus 10 synthesiser at a 50 μmol scale. Phosphoramidites (Thermo Fisher Scientific) were dissolved to 0.1 M in anhydrous acetonitrile (Merck). All reagents were used as supplied without further purification.

Oligonucleotides were cleaved and deprotected from the solid support in a mixture of aqueous ammonia (40%, 2 mL) and methylamine (33%, 2 mL) (55 °C, 50 min). Secondary deprotection of TBDMS groups was achieved with TEA•3HF (1 mL) in dimethyl sulfoxide (1 mL) (65 °C, 3 h), followed by precipitation and washing in ice-cold butanol.

Purification was performed by strong anion exchange HPLC (SAX-HPLC) using a ThermoScientific™ DNAPac™ PA 100 BioLC™ column (22 x 250 mm) on a Shimadzu LC-20AP system with a Shimadzu 20AB pump and an SPD-M40 photo diode array detector with mobile phases, A: Milli-Q™ water with sodium hydroxide (5 mM, pH 12), and B: Milli-Q™ water with sodium chloride (1 M) with sodium hydroxide (5 mM, pH 12). The concentration of mobile phase B was increased from 20-80-95% over 60 minutes with a flow rate of 5 mL/min and monitored at 260 nm. Desired fractions were collected, neutralised with dilute HCl, lyophilised, and desalted using Amicon Ultra 3k centrifugal filters.

### Secondary structure assembly and buffer conditions

RNA secondary structures were annealed in the desired buffer using an Applied Biosystems ProFlex PCR system. Oligonucleotides were heated at 85 °C for 15 min, followed by cooling at 0.1 °C/min to 20 °C, unless otherwise stated. Experiments were performed in 50 mM potassium phosphate buffers, with added KCl (50 mM) unless otherwise stated, pH adjusted to 7.00 using HCl / KOH. Unless otherwise stated, experiments were performed at 25 °C. The pH of the buffers and samples were measured with a Hanna Instruments HI11310 pH-probe.

### Ultraviolet-visible and circular dichroism spectroscopy

Oligonucleotide purity and concentration were determined using a Thermo Fisher NanoDrop One/OneC microvolume UV Spectrophotometer. 1 μL of sample in Milli-Q™ water measured at (A260/A280 ≥ 2.0) Molar extinction coefficients were calculated with the IDR OligoAnalyzer™ tool.

Circular dichroism (CD) and UV-Vis spectra were recorded in a 1 mm quartz cuvette on an Applied Photophysics Chirascan Plus CD Spectrometer. Samples (20 μM) were dissolved in Potassium Phosphate (50 mM) and Potassium Chloride (50 mM) at pH 7.01. CD spectra were obtained at 25 °C with an interval and bandwidth of 1 nm.

For melting curve titrations, RNA samples (20 mM) were prepared in 1x Tris Acetate Buffer at pH 7.03, and titrated with 1-16 equivalents of **RGGFGGRGG**. Ellipticity at 260 nm (parallel) or 295 nm (antiparallel) (63) was monitored every 0.2 °C with a temperature gradient of 0.6 °C min^-1^ from 5 – 95 °C. Data was fit to an asymmetric sigmoidal nonlinear regression to determine the melting temperature.

### Nuclear magnetic resonance (NMR) spectroscopy

^1^H NMR spectra were obtained on a Bruker Avance III 600 MHz Cryo NMR spectrometer. Peptide and oligonucleotide samples were prepared in 10% D_2_O: 90% buffer. Water suppression was achieved using excitation sculpting. Unless otherwise stated, spectra were collected at 25 °C.

Secondary structure and binding mechanism was determined by multidimensional NMR spectroscopy (HSQC, TOCSY and NOESY). Peptide:RNA proximity crosspeaks observed by NOESY were validated by saturation transfer difference (STD) NMR spectroscopy. Oligonucleotide samples were prepared to a concentration of 2 mM in 10% D_2_O: 90% buffer.

### NMR titrations

NMR titrations were performed using a constant concentration of peptide (125 μM), and titrated with 0-4 equivalents of oligonucleotide. Samples (160 μL) were prepared in 3 mm NMR tubes in 10% D_2_O: 90% buffer. A minimal concentration of 3-(trimethylsilyl)propinoic-2,2,3,3-*d*_4_ acid sodium salt was included as an internal standard. A discussion of the models and data fitting is available in Supplementary Information 1.3.

## RESULTS

### Rational design of short RNA-binding peptides derived from nucleolin

The human nucleolin RNA-binding protein is a multifunctional transcription regulator involved in ribosomal biogenesis, RNA metabolism and chromatin formation (64). Nucleolin is known interact with G-rich RNAs *in vivo* (58), with recent studies demonstrating its stabilising effect on RNA G-quuadruplex (rG4) structures (57). Nucleolin facilitates rG4 binding through its C terminal RG/RGG-rich intrinsically disordered region (IDR) (55,65).

The structural complexity and vast size of biomacromolecules make it challenging to study specific intermolecular interactions between proteins and nucleic acids, especially for weakly-associating RNAs and peptides. Using small-scale synthetic replicas of biological systems allows the probing of specific peptide-oligonucleotide interactions on a molecular level, which may lead to the future design of peptide targeting agents for therapies (66), drug delivery (67), and artificial chemical systems (68). It is estimated that up to 97% of the RNA produced in cells is non-protein-coding (69), many of which are involved in the formation of biomolecular condensates in the nucleus and cytoplasm (70,71), including the action of genetic loci called ‘enhancers’, which control the spatiotemporal; patterns of development (72,73). Unravelling the range of weak interactions of these non-coding RNAs with IDRs is key to understanding their functional roles in cellular and developmental regulation.

To elucidate the binding dynamics of IDR peptides with rG4s, three short, rationally designed peptide sequences – 1: **RGGFGGRGG** (R644-G652) (Figure 1A), 2: **RGGRGGRGG**, and 3: **LGKGGYGKV** - were derived from the Nucleolin protein. The arginine residues in peptides 1 and 2 promote affinity for the RNA phosphate backbone through electrostatic attraction and guanidinium-phosphate hydrogen bonding (74). The phenylalanine residue in peptide 1 allows for π-π stacking with the aromatic nucleobases, which is thought to be essential for nucleolin’s interactions *in vivo* (58,75). Glycine residues provide structural flexibility to the peptide backbone, allowing the peptide to conform to the RNA grooves for optimal binding (12). Peptide 3 was designed as a negative control; although it retains a 2^+^ charge, aromatic residue and glycine flexibility, it lacks the characteristic RGG repeat motif.

### RGGFGGRGG selectively binds the TERRA rG4 over the DNA counterpart

To evaluate whether **RGGFGGRGG** binds preferentially to folded RNA G-quadruplexes (rG4s), we determined the association constants between **RGGFGGRGG** and TERRA rG4 using ¹H NMR titrations performed in duplicate. NMR titrations were performed with a constant concentration of **RGGFGGRGG** (125 μM) and increasing ratios of RNA or DNA, from 0 to 2 equivalents. The change in chemical shift (Δδ) of the aromatic phenylalanine protons of the **RGGFGGRGG** peptide were monitored (Figure 2A), converted to binding isotherms (Figure 2B), and analysed using the online tool supramolecular.org (76,77), with the weighted average association constants (M^-1^) reported (Figure 2C).

**Figure 2.**
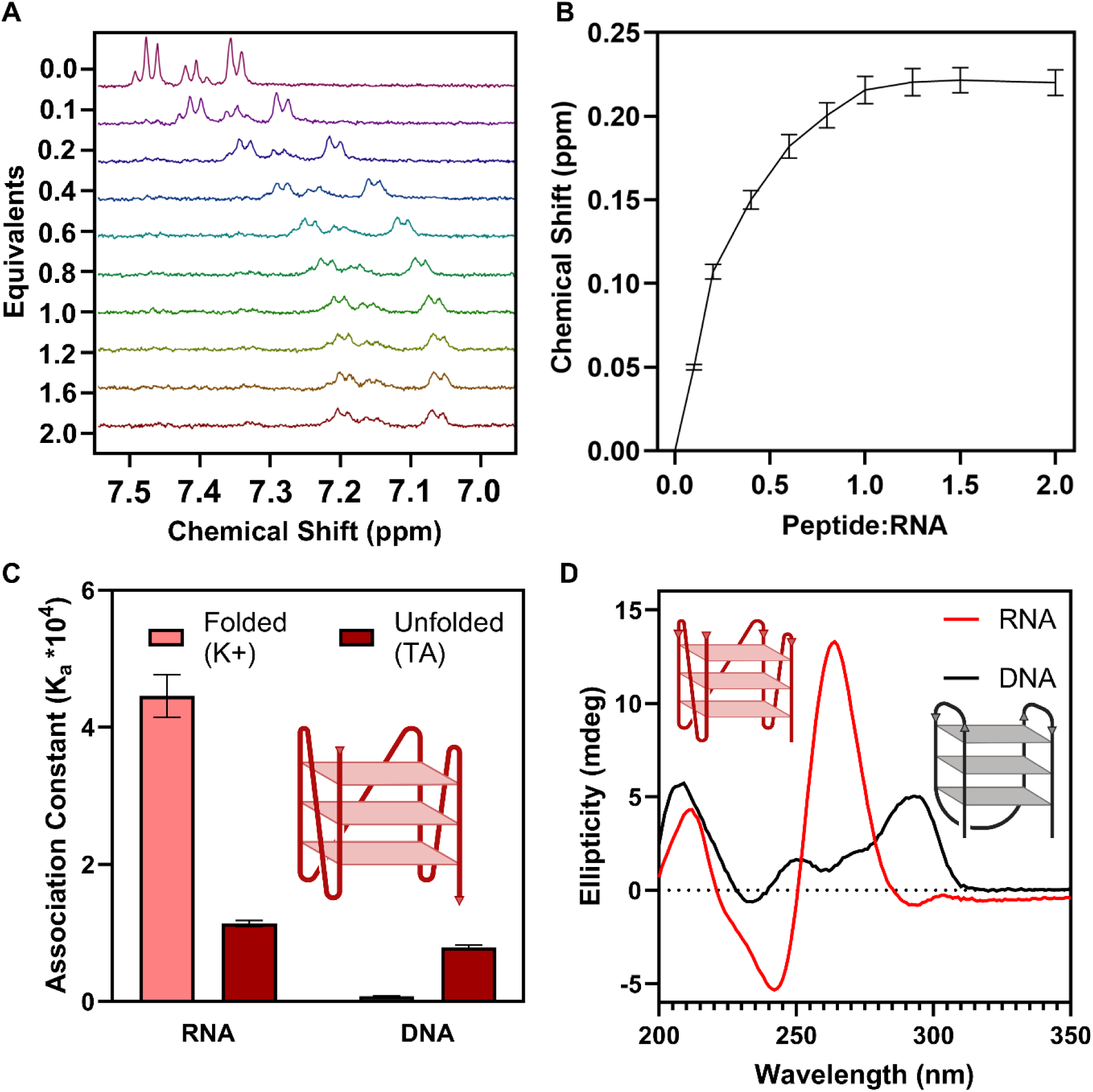
**The RGGFGGRGG peptide is selective for TERRA rG4s over a DNA counterpart, and discriminates for folded structures**. (**A):** ^1^H NMR titration data of the **RGGFGGRGG** peptide titrated with TERRA RNA (0-2 eq.). Region showed corresponds to phenylalanine aromatic protons. (**B):** Binding isotherm of the **RGGFGGRGG** aromatic phenylalanine protons upon being titrated with TERRA RNA. **(C)**: 2:1 statistical association constants determined by supramolecular.org between the peptide and G4-forming 5’-(GGGUUA)3GGG-3’ sequences: RNA vs DNA (U → T) in the folded (K^+^) or unfolded (TA) states. (**D)**: Circular dichroism spectra of TERRA RNA and DNA G4 structures. RNA forms a parallel rG4, while DNA forms a predominantly antiparallel G4, as indicated by the arrows on the schematic 5’ to 3’ backbones.

^1^H NMR titrations using the **RGGFGGRGG** peptide showed that theTERRA rG4 sequence – 5’-GGG UUA GGG UUA GGG UUA GGG-3’ – interacted with a ∼4-fold higher association constant when folded in a potassium-containing buffer, as compared to the same RNA sequence, that was unfolded in the absence of potassium (ΔΔG = –3.2 kJ mol^-1^, Figure 2C). The peptide further exhibited a ∼50-fold selectivity in terms of association constants for the stepwise 2:1 peptide-to-RNA equilibria for the folded rG4 over the DNA variant – 5’-GGG TTA GGG TTA GGG TTA GGG-3’ – (ΔΔG = –10.0 kJ mol^-1^). Isothermal Titration Calorimetry results agreed with the binding constants obtained from NMR titrations (Supplementary Information Figure 8).

We compared fitting models for the folded and unfolded states of TERRA RNA and DNA interacting with **RGGFGGRGG** (Supplementary Information Tables 2-3). In all cases, the stoichiometry was best described as two peptide molecules interacting with one nucleic acid structure. The most appropriate model for the majority of titrations was a non-cooperative statistical 2:1 host-guest model (78), which is the simplest model with the fewest forced parameters (76). This fitting model suggests that two peptide molecules bind to the TERRA rG4 independently, and that the binding of the two peptides to each nucleic acid strand is a purely statistical 2:1 process with negligible cooperativity, implying a symmetrical binding event (76,78).

We propose that the observed selectivity for **RGGFGGRGG** to RNA arises in part to the difference in folding conformation between the RNA and DNA structures. RNA exclusively adopts a parallel rG4 structure, characterised using Circular Dichroism (CD) spectroscopy by the strong positive band at 262 nm and large negative band at 245 nm (79). In contrast, the DNA G4 adopts an antiparallel topology, characterised by a positive band at 295 nm and a negative trough at 232 nm (Figure 2D) (63). These folding conformations place the tetrad sheet at a different spatial configuration, as well as orienting the loop nucleotides differently - RNA bulges to the side to form a shorter, wider ‘sandwich’ structure, while DNA protrudes at the top and bottom, to create a taller, thinner ‘chair’ structure (80). We theorise that the antiparallel G4 structures may not present an ideal binding pocket on the external backbone for the peptide to interact strongly, resulting in a decreased association constant as compared to parallel G4s.

### RGGFGGRGG templates and thermodynamically stabilises the TERRA rG4

To assess the impact of the **RGGFGGRGG** peptide on the stability of the TERRA rG4 structure, we conducted CD melting-curve titrations (9) in the absence of potassium. Hoogsteen structures - including G4s - are generally less stable than Watson-Crick helices under physiological conditions (81), and typically require precise folding conditions, such as the presence of monovalent cations for G4s. For example, rG4 structures often denature upon single point mutations, leading to a loss of function and RNA misfolding disorders (55). Selective nucleic acid binders that can influence the stability of these structures are therefore of great interest (82); however protein-inspired peptides have not featured heavily in terms of selectivity for Hoogsteen-based nucleic acid structures (68,83).

In our CD melting point assays, ellipticity at 260 nm (the characteristic wavelength of parallel G4s (63)) was monitored across a temperature gradient from 5 °C to 95 °C. The melting temperature (T_m_) was defined as the temperature where 50% of the structure had denatured. A baseline CD spectrum of TERRA RNA folded into an rG4 by 100 mM K^+^ at 5 °C was used as the ‘100% folded’ reference. All subsequent measurements were normalised to this value, to determine the percentage of folded rG4 structure upon the addition of **RGGFGGRGG**. Melting point experiments on the ‘unfolded’ TERRA, and 16 equivalents were performed in triplicate.

In the absence of salt or peptide, the melting temperature (T_m_) of the TERRA rG4 was 24.7 ± 0.1 °C. Although this measurement represents the ‘unfolded’ control, a residual CD peak at 260 nm corresponding to ∼40 % folded structure was observed, attributed to the inherit A-form helix of RNA (63) and trace sodium counterions from synthesis, which can weakly template G4 structures (63). Upon the addition of one equivalent of the **RGGFGGRGG** peptide, the proportion of folded rG4 increased to nearly 100%. UV-visible spectroscopy confirmed that the peptide folds the structure to a comparable extent as the endogenous ligand K^+^, as shown by comparable bathochromic and hypochromic shifts (Supplementary Information Figure 9). Further titration of **RGGFGGRGG** into the TERRA RNA resulted in a stepwise increase in the melting temperature of the rG4 structure. Upon the addition of 16 equivalents of peptide, the T_m_ increased to 34.0 ± 0.1 °C (ΔT_m_ = 9.3 °C, Figure 3B).

**Figure 3.**
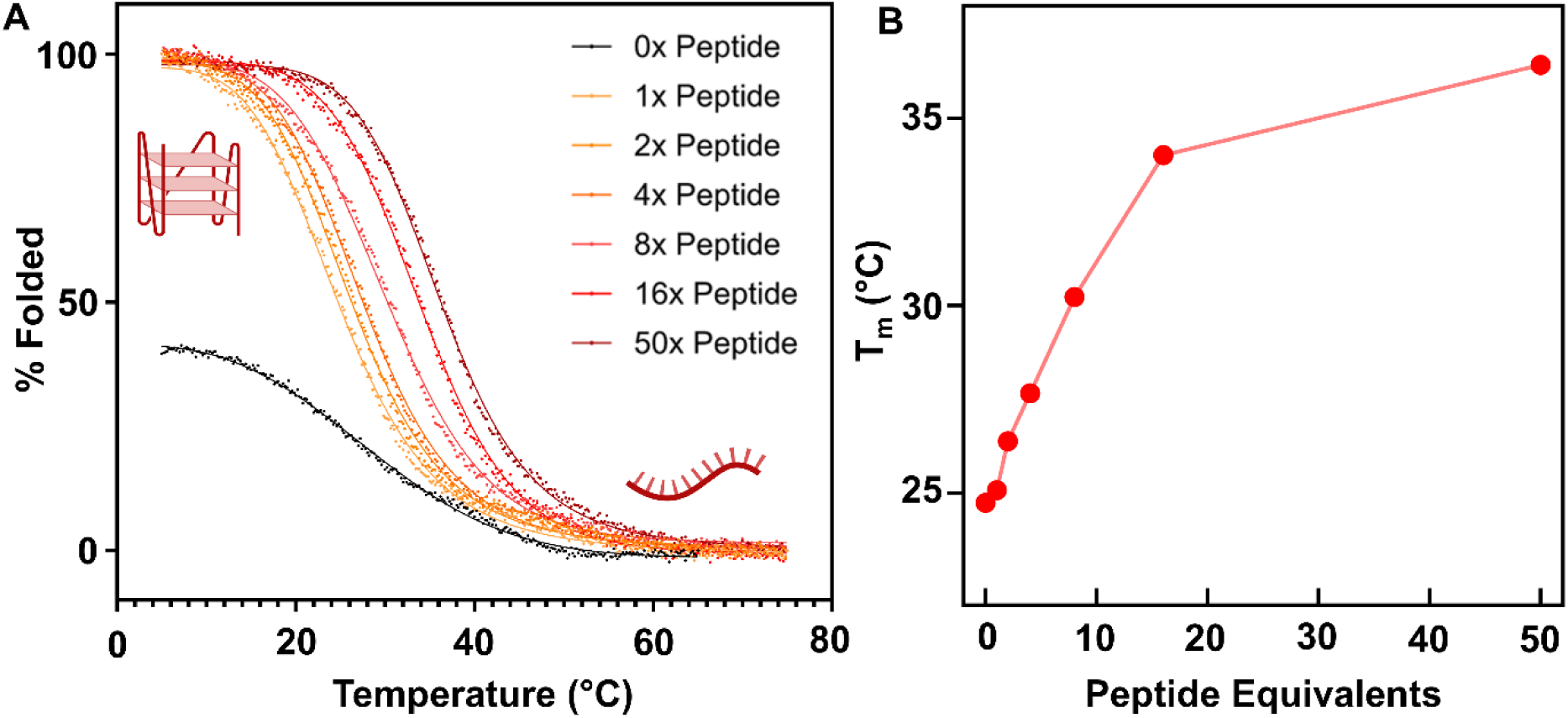
Additions of RGGFGGRGG selectively increase the thermal stability of the TERRA rG4 in the absence of potassium. **(A)**: Circular dichroism melting curves of the TERRA rG4 upon increasing equivalents of **RGGFGGRGG** (20 μM RNA, 1× tris acetate buffer at pH 7.0). Solid line is a non-linear regression best fit to the data. **(B)**: Melting temperatures (Tm) of the TERRA rG4 at each tested peptide equivalent, interpolated from the melting curves in panel A.

The stabilisation tells us that not only does **RGGFGGRGG** bind to pre-folded TERRA rG4s, but can also act as a ‘molecular chaperone’ for the TERRA rG4, promoting structure formation and increasing the thermodynamic stability even in the absence of monovalent cations. This behaviour mimics the role of cellular RNA-binding proteins that are likely essential for lncRNA structure and function *in vivo,* showing that minimalist IDR-derived peptides can replicate the structural roles of full-length proteins.

### RGGFGGRGG weakly stabilises but does not template the parallel pU27 DNA G4

RNA exclusively adopts a parallel G4 structure - no examples of antiparallel rG4s exist without modified or unnatural bases (84,85). Conversely, DNA adopts a hybrid conformation, heavily favouring antiparallel, however there are a few examples of parallel DNA G4s (86). To assess if the stabilisation of the TERRA rG4 by **RGGFGGRGG** arises primarily due to the parallel folding topology of RNA rather than sequence specificity, we conducted additional CD melting curves with a parallel DNA G4, the pU27 sequence – 5’-TGG GGA GGG TGG GGA GGG TGG GGA AGG-3’. pU27 is a G-rich tract of DNA found in the Nuclease Hypersensitive Element III1 (NHE III1) region of the c-MYC P1 promoter (87), and its parallel G4 structure has been linked to transcriptional repression of aberrant c-MYC expression, resulting in suppression of cancer proliferation (88). Studies have shown that the nucleolin protein (the source of the **RGGFGGRGG** peptide) binds and stabilises the pU27 G4 *in vivo* (58).

We first used CD spectroscopy to confirm that the pU27 sequence formed a parallel G4 in the presence of potassium (Figure 4A), in contrast to the antiparallel G4 formed by the DNA variant of TERRA. CD melting curves showed in the absence of potassium, the pU27 DNA sequence formed a partially folded parallel G4 structure (∼45% folded, T_m_ = 45.7 °C). Upon the addition of 16 equivalents of **RGGFGGRGG**, the melting point increased to 50.7 °C (ΔT_m_ = 5.0 °C), however we did not observe an increase in proportion of folded G4, suggesting that **RGGFGGRGG** modestly stabilises but does not template the pU27 G4 structure (Figure 4B). Upon the addition of 16 equivalents of the endogenous ligand K^+^, the melting point further increased to 58.6 °C (ΔT_m_ = 12.9 °C), and the parallel G4 structure was templated (Figure 4). 16 equivalents of peptide was chosen as binding simulations using supramolecular.org predicted a 90% saturation at 16 equivalents of the **RGGFGGRGG** peptide to TERRA RNA in the absence of potassium (77).

**Figure 4.**
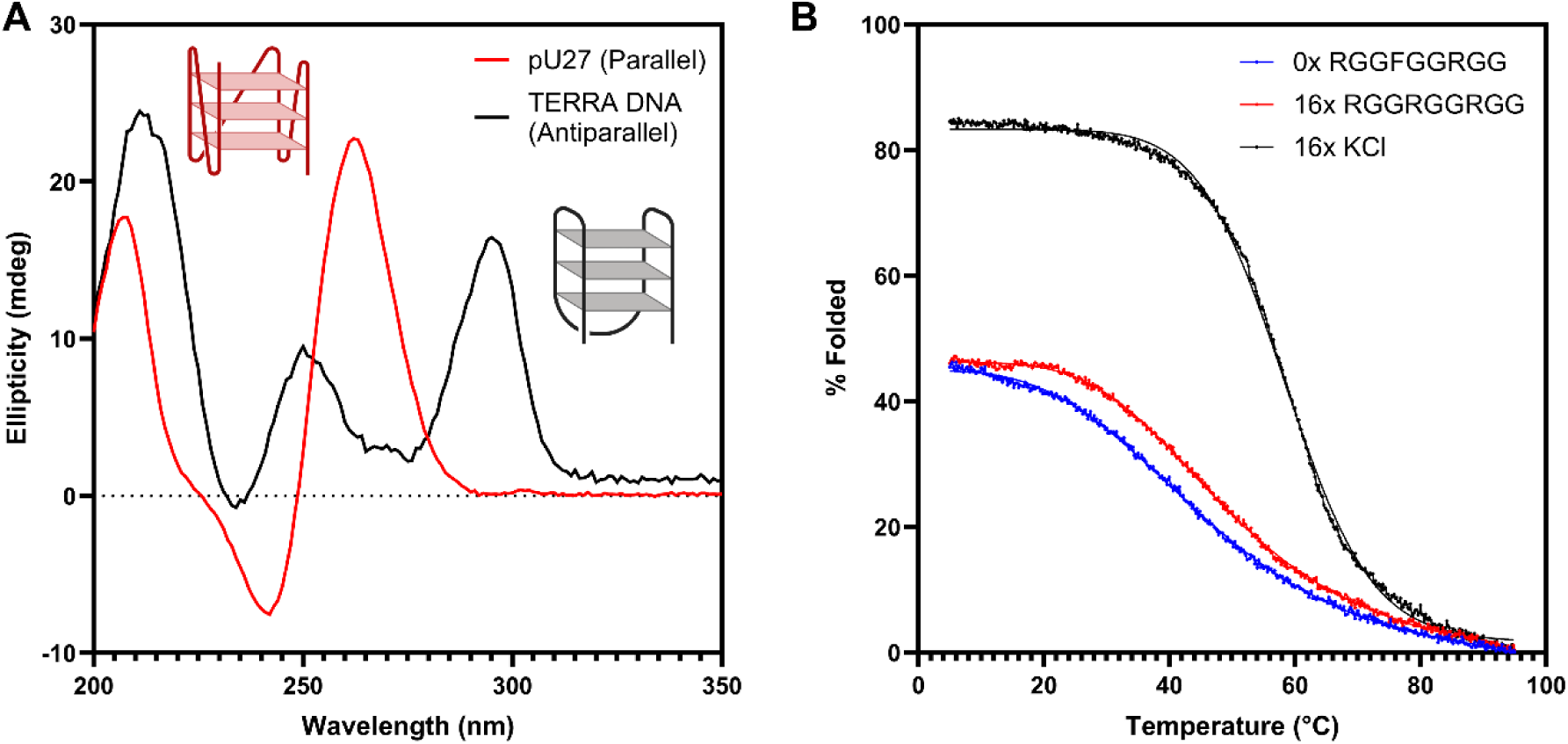
The RGGFGGRGG peptide increases the thermal stability of the parallel pU27 DNA G4 but cannot template its formation. **(A)**: Circular dichroism spectra of the parallel pU27 sequence (red), as compared to the antiparallel TERRA DNA (black) in the presence of potassium (20 μM RNA, 50 mM KCl, 50 mM potassium phosphate pH 7). **(B)**: Circular dichroism melting curves of the pU27 parallel G4 structure (blue) upon 16 equivalent additions of the **RGGFGGRGG** peptide (red) and the endogenous ligand potassium (black) (20 μM RNA, 1× tris acetate buffer at pH 7.0). Solid line is a non-linear regression best fit to the data. The **RGGFGGRGG** peptide moderately stabilises the pU27 G4 (ΔTm = 5.0 °C), whereas the endogenous ligand potassium fully templates the structure, and stabilises it to a higher degree (ΔTm = 12.9 °C).

To probe structural and sequence selectivity, we tested **RGGFGGRGG** against two additional DNA G4 sequences, the antiparallel DNA variant of the TERRA sequence – 5’-GGG TTA GGG TTA GGG TTA GGG-3’ – and an additional parallel DNA G4 sequence containing only adenines in the loop – 5’-GGG AA GGG AA GGG AA GGG-3’. Neither sequence was stabilised or templated by the **RGGFGGRGG** peptide (Supplementary Information Figure 10).

This series of CD melting curves suggests that the **RGGFGGRGG** peptide selectively stabilises the pU27 parallel DNA but cannot act as a ‘molecular chaperone’, in contrast to its chaperoning effect on the TERRA rG4. This specificity may be rationalised by the comparatively lower hydration of the DNA G4 minor groove in comparison to RNA, which may limit hydrogen bonding with the hydrophilic peptide (89), preventing it from acting as a chaperone. Additionally, the observed increase in thermodynamic stability of the parallel pU27 G4, but not for the antiparallel TERRA DNA G4, suggests that **RGGFGGRGG** interacts preferentially with parallel over antiparallel G4 structures. As **RGGFGGRGG** increases the thermal stability of the parallel pU27 sequence but not the parallel (G_3_A_2_)_n_ sequence, we propose that the interaction depends not only on structure, but also displays sequence selectivity involving loop base-identity (in this case, thymine > adenine).

### Sequence selectivity of RGGFGGRGG towards TERRA rG4

The **RGGFGGRGG** peptide both stabilises and templates the TERRA rG4. To probe its sequence selectivity for the native TERRA structure, and to determine key nucleobases required for binding, we performed additional CD melting curves on mutant TERRA sequences. From our previous results with pU27, we determined that specific loop nucleotides are significant for recruiting peptide binding, and thus stabilisation. Following this observation, we mutated the native TERRA loop sequence – UUA – to UUU and AAA, and then sequentially shortened it to two and one nucleotide (U_3_, U_2_, U_1_, A_3_, A_2_, A_1_). This approach not only demonstrates the impact of point mutations on peptide-mediated TERRA rG4 stabilisation, but also reflects the diversity of rG4 sequences found across lncRNAs. For example, a tandem repeat of (GGGA_1-3_)_n_ is representative of c-MYC associated transcripts (90,91), while (GGGU_1-3_)_n_ tandem repeats are broadly prevalent in the human genome (92). CD spectroscopy confirmed that all tested RNA sequences form rG4s (Supplementary Information Figure 11 A-F).

Upon mutation of the native TERRA tandem repeat (GGG UUA)_n_ to a loop containing only uracil (G_3_U_3_)_n_, there was a reduction in the observed stabilisation in melting point (ΔT_m_) to ∼4 °C. Although there was still minor stabilisation observed, the **RGGFGGRGG** peptide no longer induced complete formation of the rG4 structure, and the extent to which the G4 structure was folded at 25 °C was no longer at a comparable magnitude to potassium. (Table 1, Supplementary Information Figure 12 A-F). Reductions of the length of the uracil loop length progressively decreased stabilisation (Table 1). When the loop sequence was instead mutated to only contain adenine (G_3_A_3-1_)_n_, **RGGFGGRGG** lost all stabilising ability, and instead a slight destabilisation of the rG4 was observed (ΔT_m_ = –2 °C). In contrast, additions of the endogenous ligand K^+^ increased the stability of all tested mutant sequences, and induced formation of the rG4 structure (Table 1, Supplementary Information Figure 12 A-F).

**Table 1.** RGGFGGRGG selectively stabilises the native TERRA rG4 structure over loop mutation sequences, whereas the endogenous ligand K^+^ stabilises all rG4 structures. Melting temperatures (T_m_) were determined by CD melting curves, with uncertainty derived from nonlinear fitting error. The change in melting temperature (ΔT) is given relative to the control in the absence of monovalent cations. Each sample was measured with either 16 equivalents of **RGGFGGRGG** or the endogenous ligand, K^+^.

|  | Control | RGGFGGRGG (16x) |  | KCl (16x) |  |
| --- | --- | --- | --- | --- | --- |
| Sequence (5'-3') | $T_m$ (°C) | $T_m$ (°C) | $\Delta T$ (°C) | $T_m$ (°C) | $\Delta T$ (°C) |
| **(GGGUUA)<sub>n</sub>**<br>**(TERRA)** | **24.7 ± 0.1 °C** | **34.0 ± 0.1 °C** | **9.3** | **46.3 ± 0.2 °C** | **21.5** |
| (G<sub>3</sub>U<sub>3</sub>)<sub>n</sub> | 31.3 ± 0.3 °C | 35.4 ± 0.2 °C | 4.1 | 45.3 ± 0.2 °C | 14.0 |
| (G<sub>3</sub>U<sub>2</sub>)<sub>n</sub> | 36.0 ± 0.1 °C | 40.9 ± 0.2 °C | 4.8 | 54.4 ± 0.1 °C | 18.4 |
| (G<sub>3</sub>U<sub>1</sub>)<sub>n</sub> | 50.9 ± 0.2 °C | 52.5 ± 0.2 °C | 2.4 | 61.8 ± 0.2 °C | 10.9 |
| (G<sub>3</sub>A<sub>3</sub>)<sub>n</sub> | 43.9 ± 0.2 °C | 41.5 ± 0.1 °C | -2.4 | 58.7 ± 0.6 °C | 14.9 |
| (G<sub>3</sub>A<sub>2</sub>)<sub>n</sub> | 42.7 ± 0.2 °C | 36.9 ± 1.4 °C | -5.8 | 45.3 ± 0.2 °C | 2.6 |
| (G<sub>3</sub>A<sub>1</sub>)<sub>n</sub> | 52.7 ± 0.6 °C | 52.5 ± 0.1 °C | -0.3 | 59.6 ± 0.2 °C | 6.9 |

These results suggest that uracil is an essential loop residue for recruiting peptide binding, leading to stabilisation of the TERRA rG4 structure, whereas adenine loops are unfavourable for **RGGFGGRGG** interactions, mimicking the conclusions of the pU27 screens, where thymine was preferable over adenine. As specific G4 loop base identity is significant in recruiting peptide interaction, it is evident that electrostatics alone are not responsible for these RNA-peptide binding interactions (93). Notably, no tested permutation of loop sequence was stabilised by the peptide to the extent of the native TERRA sequence, highlighting the significance of evolved sequence selectivity of the **RGGFGGRGG** peptide towards the native TERRA rG4.

### Phenylalanine is essential for peptide-mediated TERRA rG4 stabilisation

Having established that the parallel TERRA rG4 structure and uracil loop nucleotides are essential for **RGGFGGRGG** interaction and stabilisation, we next mutated the peptide sequence to determine the contribution of individual amino acid residues. Biological studies have shown that nucleolin can stabilise large tracks of guanine-rich DNA. Upon mutation of nucleolin phenylalanine residues, there was a loss of stabilisation (58). To investigate the importance of phenylalanine in our system, we introduced a point mutation (4F→R) to afford the mutated peptide **RGGRGGRGG**. This mutation removes the ability of the peptide to π-π stack with the guanine tetrad of the rG4 structure while retaining its electrostatic attraction to the phosphate backbone via arginine residues.

We conducted additional CD melting curves with 16 equivalents of **RGGRGGRGG** added to TERRA RNA, comparing the thermodynamic stabilisation to the native peptide **RGGFGGRGG** (ΔT_m_ = 9.3 °C). Our study showed that upon the addition of 16 equivalents of the mutated peptide **RGGRGGRGG**, there was no stabilisation of the TERRA rG4 structure, instead a minor destabilisation was observed (Supplementary Information Figure 13A).

Compared to the native peptide, the lack of stabilising effect suggests that although the mutated **RGGRGGRGG** peptide may still interact with the TERRA rG4 structure, the phenylalanine residue is essential for π-π stacking within the guanine tetrad, providing structural stabilisation.

We tested an additional non-interacting peptide sequence (**LGKGGYGKV**) retaining 2^+^ charge. This peptide substitutes the positively charged arginine for lysine, the aromatic phenylalanine for tyrosine, and disrupts the canonical RGG RNA-binding motif. **LGKGGYGKV** had no stabilising or destabilising effect on the melting point or folding capabilities of the rG4structure, suggesting a complete lack of interaction (Supplementary Information Figure 13B). This further suggests that interaction and stabilisation of the TERRA rG4 by **RGGFGGRGG** is not driven exclusively by electrostatics – both the arginine and phenylalanine residues, as well as the RGG motif are essential.

### Structural model of RGGFGGRGG binding to TERRA rG4

Our mutation studies identified uracil loop residues in TERRA, and phenylalanine and arginine residues in **RGGFGGRGG** as essential for stabilisation of the TERRA rG4 structure. To confirm the significance of these components and to model a mechanism of interaction, we employed multidimensional NMR spectroscopy to probe through-space interactions between **RGGFGGRGG** and TERRA in the presence of potassium ions (94).

Using ^1^H-^1^H Nuclear Overhauser Effect (NOESY) NMR spectroscopy, we observed strong NOE crosspeaks between the phenylalanine aromatic protons (δ 7.2 ppm) and the guanine tetrad imino-exchange protons (δ 10.5-11.5 ppm, Figure 5A), as well as additional crosspeaks with guanine base protons (δ 7.6 ppm, Supplementary Information Figure 14). These crosspeaks suggest the phenylalanine is π-π stacking with the guanine tetrad on one extremity of the rG4 structure. Additional NOEs were observed between the phenylalanine β-CH_2_ protons (δ2.5 – 2.7 ppm), guanine sugar protons (δ5.8 ppm), and guanine base protons (δ 7.6 ppm), providing further support for an interaction between phenylalanine and guanine, as our mutation studies suggested (Supplementary Information Figure 14).

**Figure 5.**
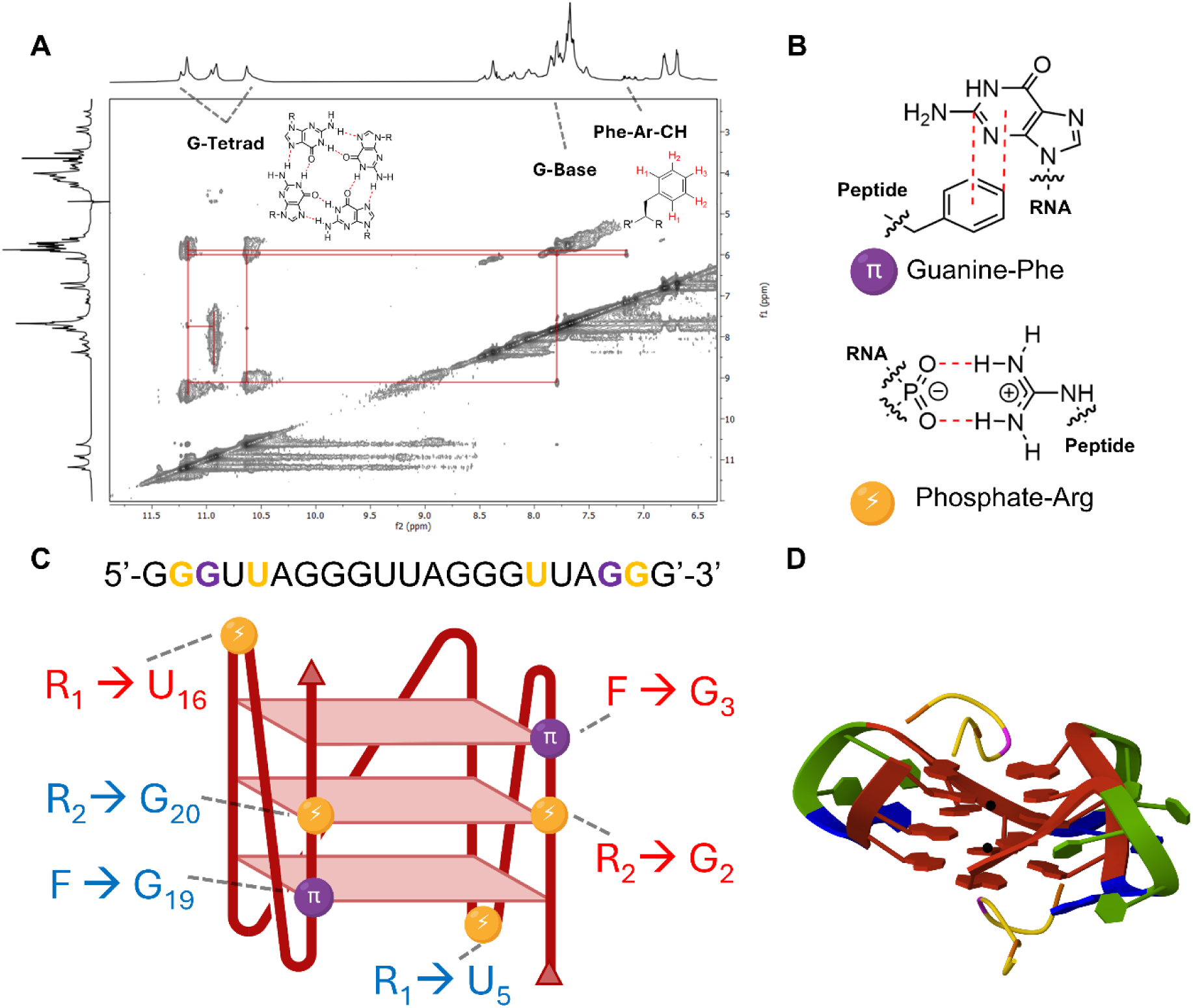
Proposed binding interactions between the RGGFGGRGG peptide and TERRA rGT4 in the presence of potassium. (A): ^1^H-^1^H NOESY NMR data showing crosspeaks between phenylalanine aromatic protons of **RGGFGGRGG** and the guanine-tetrad base protons of the TERRA rG4. **(B):** Molecular representation of phosphate-arginine (guanidinium) and guanine-phenylalanine interactions present in the host-guest complex. (**C):** Schematic representation of the TERRA rG4 structure, showing proposed interaction locations as derived from 2D NMR spectroscopy. Yellow dots represent proposed phosphate-guanidinium interactions. Purple dots represent proposed guanine-phenylalanine π-stacking. Interaction locations between the RNA and peptide phenylalanine (F) and arginine (R) are colour coded for the two binding events. **(D):** AlphaFold 3 computational prediction validating our proposed model. Two **RGGFGGRGG** peptides bind symmetrically to the top and bottom of the TERRA rG4 structure. The guanine tetrad (red) is being templated by potassium ions (black). Phenylalanine (pink) is π-stacking with guanine, arginine (orange) is interacting with the phosphate backbone associated with one guanine and one uracil (blue). Adenine (green) does not have a detected interaction with the peptide.

We also observed NOE crosspeaks between β-CH_2_ protons of arginine (δ 1.7 ppm) and both the 2’/3’ ribose protons associated with one uracil and one guanine (δ 3.0 – 4.5 ppm, Supplementary Information Figure 15), suggesting guanidinium-phosphate hydrogen bonding and electrostatic interactions on the external phosphate backbone of the G4 structure (74). The precise assignment of these protons in the RNA sequence are moderately ambiguous, as it is not possible to determine which specific uracil/guanine is interacting, as we cannot resolve individual bases without isotope labelling (95). Saturation transfer difference (STD) NMR spectroscopy epitope mapping mirrored NOESY structural data, showing strong proximities between the peptide with guanine bases, and guanine / uracil sugars (Supplementary Information Figure 16).

Combining the mutation studies and NMR data with our 2:1 peptide-to-RNA symmetrical stoichiometry from NMR titrations, we propose a model where the peptide binds symmetrically to both the top and bottom of the G4 structure. In both binding events, we predict three interaction sites - two arginine residues forming electrostatic and hydrogen-bonds with the TERRA phosphate backbone associated with one guanine and one uracil (Figure 5B, bottom) (74), and one phenylalanine π-π stacking with the guanine tetrad (Figure 5B, top).

To validate this model, we generated a computational prediction using AlphaFold 3 (96). We first modelled the native TERRA sequence with 2 potassium ions, which produced a correctly folded parallel rG4, validating AlphaFold 3 as capable of predicting non-canonical RNA structures. After, we added two copies of the **RGGFGGRGG** peptide, and the proposed model agreed with our mechanism following simulation (Figure 5D). The model places the peptides symmetrically at the top and bottom of the G4 structure, with the same three non-covalent interactions between each peptide and the RNA. In the AlphaFold model, phenylalanine (F_4_) π stacks with guanine (1F_4_ →G_3_, 2F_4_ →G_19_), the first arginine (R_1_) in each peptide interacts with the phosphate backbone associated with uracil (1R_1_ →U_5_, 2R_1_ →U_16_), and the second arginine (R_7_) in each peptide interacts with the phosphate backbone associated with guanine (1R_7_ →G_2_ 2R_7_ →G_20_).

Together, the experimental and computational data combined provides strong reasoning for the proposed interaction mechanism for the minimal peptide **RGGFGGRGG** interacting with a short model of human telomeric RNA. This mechanism provides insight into the harmonic cooperation between aromatic and cationic residues to stabilise the TERRA rG4 structure through a combination of π-π stacking and guanidinium mediated electrostatic templation.

## DISCUSSION

RNA G-quadruplex (rG4) structures in long non-coding RNAs (lncRNAs) are emerging as areas of therapeutic interest (97). Their structures *in vivo* are shaped by not only physicochemical conditions – ionic strength, monovalent cation concentration and pH – but also the macromolecular environment – including interactions with RNA binding proteins (RBPs) (55,56,58,98) or other nucleic acids (99), and macromolecular crowding (59), features that are often overexpressed in proteins containing intrinsically disordered regions (IDRs), which typically bind RNA structures (18–20, 35–37).

Currently available G4 stabilisers are limited to small molecule ligands – TMPyP4, BRACO-19, PhenDC3 & PDS (100–103). While these ligands bind to and stabilise G4 structures with high affinity, they often show limited discrimination between DNA and RNA G4s (102). Furthermore, they tend to bind and/or stabilise a diverse range of G4 structures with little to no sequence selectivity,and can also bind other nucleic acid secondary structures (e.g., i-motifs, triplexes, and even duplex or single stranded regions) (82,104). This promiscuity lacks the discrimination and biocompatibility required in cellular applications.

G4 structures are stabilised through two main mechanisms: *endo*-and *exo*-stabilization. Monovalent cations stabilise G4s via *endo*-stabilisation, coordinating with the central guanine tetrad (9) and promoting self-complementary Hoogsteen hydrogen bonding. Small molecule stabilisers (e.g., TMPyP4) are thought to work through both *endo-* and *exo-*stabilisation via intercalation and end-stacking terminal tetrads, impeding unfolding (100). Given our NMR data, we propose that the stabilisation of rG4s by **RGGFGGRGG** relies on *exo*-stabilisation through interactions with the external phosphate backbone of the RNA structure, and π-π stacking with the terminal tetrads. These external interactions are the main mechanism for cellular protein stabilisation of rG4s in biological systems (29, 55, 87–89).

Peptide-based targeting agents and therapeutics offer orthogonal interactions, leading to inherit specificity and the ability to rationally design sequences to form transient, tunable interactions with biomolecules, while being extremely viable for scalable synthesis and remaining biocompatible. Clinically validated peptide-based drugs (e.g., semaglutide) provide examples of macromolecular interactions being used with therapeutic and commercial success (105). The **RGGFGGRGG** peptide demonstrates that even a minimalistic 9-mer peptide can provide selective recognition of complex RNA structures *in vitro* while retaining biological function, using only a small set of highly conserved amino acid residues (106). This minimalistic design could be expanded into a library of short RGG-based peptides by systematically varying glycine content to modulate structural flexibility and performing orthogonal screens to probe, stabilise, or inhibit lncRNA secondary structures. Further, alternate peptide sequences can be rationally derived from IDRs of RBPs such as FMRP, hnRNP A1, FUS or TDP-43 (55,107–109), to target their native RNA substrates and mimic functions such as structural stabilisation, oncogene regulation, and transcriptional / translational regulation.

While the **RGGFGGRGG** peptide is a promising minimalist model for selective TERRA rG4 stabilisation, there are several limitations to be considered before this approach can be extended to biological systems. The relatively modest binding affinity requires a high stoichiometric ratio of peptide to reach saturation, which is a problem *in vivo* due to low RNA concentration and high competition with RBPs (41). The flexibility afforded by the glycines is useful as a screening tool to maximise the change of an interaction, but it reduces the overall binding strength of the peptide. Additionally, while the short length of the peptide is a significant advantage in supramolecular and physical chemistry applications, it may undergo rapid enzymatic degradation *in vivo* (110). These drawbacks can be partially mitigated by reducing the affinity of cellular proteases to the peptide by making unnatural modifications to the sequence, such as arginine methylation or N/C terminal capping, which can slow degradation (111,112). Future advances in delivery devices and targeting strategies may expand the uses of minimalist peptides to compete in the cellular environment.

In summary, this study presents the **RGGFGGRGG** peptide, derived from the IDR of the RNA-binding protein nucleolin, as a minimalist molecular chaperone that is selective for the human telomeric repeat containing lncRNA (TERRA) rG4 structure. This peptide works in the absence of monovalent cations and displays selectivity for nucleic acid identity (RNA over DNA), topology (parallel G4 over antiparallel G4) and specific loop base identity (uracil over adenine). Multidimensional NMR and computational modelling agreed on a symmetrical 2:1 peptide:RNA interaction mechanism, in which phenylalanine π-π stacks with guanine tetrads, while arginine residues engage in electrostatic and hydrogen bonding with the phosphate backbone near uracil and guanine residues (53–55, 62). Together, these conclusions rationalise our observed selectivity of **RGGFGGRGG** for TERRA, offering a stepping stone for mimicking RBP functions on lncRNA regulation.

## DATA AVAILABILITY

This research did not generate any sensitive datasets. Unaltered raw data is available either in the Supplementary Information or is available upon request. NMR titration data is available via a GitHub repository: https://github.com/pallithordarson/RNA_quadruplex_data

## SUPPLEMENTARY DATA

The supplementary data includes characterization by HPLC, LCMS, NMR and CD, experimental procedures, raw thermographs and multidimensional NMR.

## ACKNOWLEDGEMENTS

The authors would like to acknowledge the Mark Wainwright Analytical Centre at the University of New South Wales for providing access to instruments at the NMR Facility, Structural Biology Facility and Bioanalytical Mass Spectra Facility. They also thank the UNSW RNA Institute – in particular Dr. Hsiu Lin Li – for providing the oligonucleotides used in this project and Dr. Susannah L. Brown for mass spectra analysis. The authors thank Prof. (E. W.) Bert Meijer, Dr. Julian Huppert and Mr. Christopher Pracey for helpful conversations. L.B.C. is grateful to the Australian Research Council and UNSW for a PhD scholarship. The authors acknowledge that Biorender.com was used to create the graphics used in this manuscript.

## REFERENCES

1. Bernat, V. and Disney, M.D. RNA Structures as Mediators of Neurological Diseases and as Drug Targets.

2. Devi, G., Zhou, Y., Zhong, Z., Toh, D.-F.K. and Chen, G. (2015) RNA triplexes: from structural principles to biological and biotech applications. WIREs RNA, 6, 111–128.

3. Timsit, Y. and Moras, D. (1996) Cruciform structures and functions. Quarterly Reviews of Biophysics, 29, 279–307.

4. Cox, L., Bai, C., Platnich, C.M. and Rizzuto, F.J. (2024) Divergent Polymer Superstructures from Protonated Poly(adenine) DNA and RNA. Biomacromolecules, 25, 3163–3168.

5. King, J.J., Irving, K.L., Evans, C.W., Chikhale, R.V., Becker, R., Morris, C.J., Peña Martinez, C.D., Schofield, P., Christ, D., Hurley, L.H. et al. (2020) DNA G-Quadruplex and i-Motif Structure Formation Is Interdependent in Human Cells. Journal of the American Chemical Society, 142, 20600–20604.

6. Bertuch, A.A. (2016) The molecular genetics of the telomere biology disorders. RNA Biology, 13, 696–706.

7. La Spada, A.R., Paulson, H.L. and Fischbeck, K.H. (1994) Trinucleotide repeat expansion in neurological disease. Annals of Neurology, 36, 814–822.

8. Kishtagari, A. and Watts, J. (2017) Biological and clinical implications of telomere dysfunction in myeloid malignancies. Therapeutic Advances in Hematology, 8, 317–326.

9. Huppert, J.L. (2008) Four-stranded nucleic acids: structure, function and targeting of G-quadruplexes. Chemical Society Reviews, 37, 1375–1384.

10. Simone, R., Fratta, P., Neidle, S., Parkinson, G.N. and Isaacs, A.M. (2015) G-quadruplexes: Emerging roles in neurodegenerative diseases and the non-coding transcriptome. FEBS Letters, 589, 1653–1668.

11. Lyu, K., Chow, E.Y.-C., Mou, X., Chan, T.-F. and Kwok, Chun K. (2021) RNA G-quadruplexes (rG4s): genomics and biological functions. Nucleic Acids Research, 49, 5426–5450.

12. Chowdhury, M.N. and Jin, H. (2023) The RGG motif proteins: Interactions, functions, and regulations. WIREs RNA, 14, e1748.

13. Lipps, H.J. and Rhodes, D. G-quadruplex structures: in vivo evidence and function.

14. Montero, J.J., López de Silanes, I., Graña, O. and Blasco, M.A. (2016) Telomeric RNAs are essential to maintain telomeres. Nature Communications, 7, 12534.

15. Luke, B. and Lingner, J. (2009) TERRA: telomeric repeat-containing RNA. The EMBO Journal, 28, 2503–2510-2510.

16. Chebly, A.A.-O., Ropio, J., Baldasseroni, L., Prochazkova-Carlotti, M., Idrissi, Y., Ferrer, J., Farra, C., Beylot-Barry, M., Merlio, J.A.-O. and Chevret, E. Telomeric Repeat-Containing RNA (TERRA): A Review of the Literature and First Assessment in Cutaneous T-Cell Lymphomas. LID - 10.3390/genes13030539 [doi] LID - 539.

17. Uversky, V.N. (2013) A decade and a half of protein intrinsic disorder: Biology still waits for physics. Protein Science, 22, 693–724.

18. Uversky, V.N. (2016) Dancing protein clouds: the strange biology and chaotic physics of intrinsically disordered proteins. Journal of Biological Chemistry, 291, 6681–6688.

19. Uversky, V.N. (2019) Intrinsically disordered proteins and their “mysterious” (meta)physics. Frontiers in Physics, 7, article 10.

20. Kulkarni, P. and Uversky, V.N. (2018) Intrinsically disordered proteins: the dark horse of the dark proteome. Proteomics, 18, article 1800061.

21. Minezaki, Y., Homma, K., Kinjo, A.R. and Nishikawa, K. (2006) Human transcription factors contain a high fraction of intrinsically disordered regions essential for transcriptional regulation. Journal of Molecular Biology, 359, 1137–1149.

22. Korneta, I. and Bujnicki, J.M. (2012) Intrinsic disorder in the human spliceosomal proteome. PLOS Computational Biology, 8, e1002641.

23. Tantos, A., Han, K.-H. and Tompa, P. (2012) Intrinsic disorder in cell signaling and gene transcription. Molecular and Cellular Endocrinology, 348, 457–465.

24. Chen, W. and Moore, M.J. (2014) The spliceosome: disorder and dynamics defined. Current Opinion in Structural Biology, 24, 141–149.

25. Staby, L., O’Shea, C., Willemoës, M., Theisen, F., Kragelund, B.B. and Skriver, K. (2017) Eukaryotic transcription factors: paradigms of protein intrinsic disorder. Biochemical Journal, 474, 2509–2532.

26. Niklas, K.J., Dunker, A.K. and Yruela, I. (2018) The evolutionary origins of cell type diversification and the role of intrinsically disordered proteins. Journal of Experimental Botany, 69, 1437–1446.

27. Liu, J., Perumal, N.B., Oldfield, C.J., Su, E.W., Uversky, V.N. and Dunker, A.K. (2006) Intrinsic disorder in transcription factors. Biochemistry, 45, 6873–6888.

28. Lazar, T., Schad, E., Szabo, B., Horvath, T., Meszaros, A., Tompa, P. and Tantos, A. (2016) Intrinsic protein disorder in histone lysine methylation. Biology Direct, 11, 30.

29. Wright, P.E. and Dyson, H.J. (2015) Intrinsically disordered proteins in cellular signalling and regulation. Nature Reviews Molecular Cell Biology, 16, 18–29.

30. Hahn, S. (2018) Phase separation, protein disorder, and enhancer function. Cell, 175, 1723–1725.

31. Peng, Z., Mizianty, M.J., Xue, B., Kurgan, L. and Uversky, V.N. (2012) More than just tails: intrinsic disorder in histone proteins. Molecular BioSystems, 8, 1886–1901.

32. Watson, M. and Stott, K. (2019) Disordered domains in chromatin-binding proteins. Essays in Biochemistry, 63, 147–156.

33. Balcerak, A., Trebinska-Stryjewska, A., Konopinski, R., Wakula, M. and Grzybowska, E.A. (2019) RNA–protein interactions: disorder, moonlighting and junk contribute to eukaryotic complexity. Open Biology, 9, article 190096.

34. Quintero-Cadena, P., Lenstra, T.L. and Sternberg, P.W. (2020) RNA Pol II length and disorder enable cooperative scaling of transcriptional bursting. Molecular Cell, 79, 207–220.

35. Musselman, C.A. and Kutateladze, T.G. (2021) Characterization of functional disordered regions within chromatin-associated proteins. iScience, 24, article 102070.

36. Zeke, A., Schád, É., Horváth, T., Abukhairan, R., Szabó, B. and Tantos, A. (2022) Deep structural insights into RNA-binding disordered protein regions. WIREs RNA, 13, e1714.

37. Polymenidou, M. (2018) The RNA face of phase separation. Science, 360, 859–860.

38. Roden, C. and Gladfelter, A.S. (2021) RNA contributions to the form and function of biomolecular condensates. Nat Rev Mol Cell Biol, 22, 183–195.

39. Marcotte, E.M. and Tsechansky, M. (2009) Disorder, promiscuity, and toxic partnerships. Cell, 138, 16–18.

40. Cumberworth, A., Lamour, G., Babu, M.M. and Gsponer, J. (2013) Promiscuity as a functional trait: intrinsically disordered regions as central players of interactomes. Biochemical Journal, 454, 361–369.

41. Davidovich, C., Zheng, L., Goodrich, K.J. and Cech, T.R. (2013) Promiscuous RNA binding by Polycomb repressive complex 2. Nature Structural & Molecular Biology, 20, 1250–1257.

42. Davidovich, C. and Cech, T.R. (2015) The recruitment of chromatin modifiers by long noncoding RNAs: lessons from PRC2. RNA, 21, 2007–2022.

43. Davidovich, C., Wang, X., Cifuentes-Rojas, C., Goodrich, Karen J., Gooding, Anne R., Lee, Jeannie T. and Cech, Thomas R. (2015) Toward a consensus on the binding specificity and promiscuity of PRC2 for RNA. Molecular Cell, 57, 552–558.

44. Cajigas, I., Leib, D.E., Cochrane, J., Luo, H., Swyter, K.R., Chen, S., Clark, B.S., Thompson, J., Yates, J.R., III, Kingston, R.E., et al. (2015) Evf2 *lncRNA/BRG1/DLX1* interactions reveal RNA-dependent inhibition of chromatin remodeling. Development, 142, 2641–2652.

45. Frege, T. and Uversky, V.N. (2015) Intrinsically disordered proteins in the nucleus of human cells. Biochemistry and Biophysics Reports, 1, 33–51.

46. Protter, D.S.W., Rao, B.S., Van Treeck, B., Lin, Y., Mizoue, L., Rosen, M.K. and Parker, R. (2018) Intrinsically disordered regions can contribute promiscuous interactions to RNP granule assembly. Cell Reports, 22, 1401–1412.

47. Brodsky, S., Jana, T., Mittelman, K., Chapal, M., Kumar, D.K., Carmi, M. and Barkai, N. (2020) Intrinsically disordered regions direct transcription factor *in vivo* binding specificity. Molecular Cell, 79, 459–471.

48. Chakrabarti, P. and Chakravarty, D. (2022) Intrinsically disordered proteins/regions and insight into their biomolecular interactions. Biophysical Chemistry, 283, article 106769.

49. Papagiannoula, A., Vedel, I.M., Motzny, K., Tengo, M., Saiti, A. and Milles, S. (2025) Promiscuous and multivalent interactions between Eps15 and partner protein Dab2 generate a complex interaction network. Nature Communications, 16, article 7783.

50. Niklas, K.J., Bondos, S.E., Dunker, A.K. and Newman, S.A. (2015) Rethinking gene regulatory networks in light of alternative splicing, intrinsically disordered protein domains, and post-translational modifications. Frontiers in Cell and Developmental Biology, 3, article 8.

51. Macossay-Castillo, M., Marvelli, G., Guharoy, M., Jain, A., Kihara, D., Tompa, P. and Wodak, S.J. (2019) The balancing act of intrinsically disordered proteins: enabling functional diversity while minimizing promiscuity. Journal of Molecular Biology, 431, 1650–1670.

52. Gsponer, J., Futschik, M.E., Teichmann, S.A. and Babu, M.M. (2008) Tight regulation of unstructured proteins: from transcript synthesis to protein degradation. Science, 322, 1365–1368.

53. Vavouri, T., Semple, J.I., Garcia-Verdugo, R. and Lehner, B. (2009) Intrinsic protein disorder and interaction promiscuity are widely associated with dosage sensitivity. Cell, 138, 198–208.

54. Jain, A. and Vale, R.D. (2017) RNA phase transitions in repeat expansion disorders. Nature, 546, 243–247.

55. Ozdilek, B.A., Thompson, V.F., Ahmed, N.S., White, C.I., Batey, R.T. and Schwartz, J.C. (2017) Intrinsically disordered RGG/RG domains mediate degenerate specificity in RNA binding. Nucleic Acids Research, 45, 7984–7996.

56. Phan, A.T., Kuryavyi, V., Darnell, J.C., Serganov, A., Majumdar, A., Ilin, S., Raslin, T., Polonskaia, A., Chen, C., Clain, D. et al. (2011) Structure-function studies of FMRP RGG peptide recognition of an RNA duplex-quadruplex junction. Nature Structural & Molecular Biology, 18, 796–804.

57. Masuzawa, T. and Oyoshi, T. (2020) Roles of the RGG Domain and RNA Recognition Motif of Nucleolin in G-Quadruplex Stabilization. ACS Omega, 5, 5202–5208.

58. González, V., Guo, K., Hurley, L. and Sun, D. (2009) Identification and Characterization of Nucleolin as a c-<em>myc</em> G-quadruplex-binding Protein *. Journal of Biological Chemistry, 284, 23622–23635.

59. Moller, A.L., Middleton, I.A., Maynard, G.E., Cox, L.B., Wang, A., Li, H.L. and Thordarson, P. (2025) Discrimination between Purine and Pyrimidine-Rich RNA in Liquid–Liquid Phase-Separated Condensates with Cationic Peptides and the Effect of Artificial Crowding Agents. Biomacromolecules, 26, 470–479.

60. Mei, Y., Deng, Z., Vladimirova, O., Gulve, N., Johnson, F.B., Drosopoulos, W.C., Schildkraut, C.L. and Lieberman, P.M. (2021) TERRA G-quadruplex RNA interaction with TRF2 GAR domain is required for telomere integrity. Scientific Reports, 11, 3509.

61. Castello, A., Fischer, B., Hentze, M.W. and Preiss, T. (2013) RNA-binding proteins in Mendelian disease. Trends in Genetics, 29, 318–327.

62. Meyer, K., Kirchner, M., Uyar, B., Cheng, J.-Y., Russo, G., Hernandez-Miranda, L.R., Szymborska, A., Zauber, H., Rudolph, I.-M., Willnow, T.E. et al. (2018) Mutations in disordered regions can cause disease by creating dileucine motifs. Cell, 175, 239–253.

63. del Villar-Guerra, R., Trent, J.O. and Chaires, J.B. (2018) G-Quadruplex Secondary Structure Obtained from Circular Dichroism Spectroscopy. Angewandte Chemie International Edition, 57, 7171–7175.

64. Mongelard, F. and Bouvet, P. (2007) Nucleolin: a multiFACeTed protein. Trends in Cell Biology, 17, 80–86.

65. Thandapani, P., O’Connor, T.R., Bailey Timothy L. and Richard, S. (2013) Defining the RGG/RG Motif. Molecular Cell, 50, 613–623.

66. Scionti, F., Juli, G., Rocca, R., Polerà, N., Nadai, M., Grillone, K., Caracciolo, D., Riillo, C., Altomare, E., Ascrizzi, S. et al. (2023) TERRA G-quadruplex stabilization as a new therapeutic strategy for multiple myeloma. Journal of Experimental & Clinical Cancer Research, 42, 71.

67. Walker, M.J. and Varani, G. (2019) In Hargrove, A. E. (ed.), Methods in Enzymology. Academic Press, Vol. 623, pp. 339–372.

68. Shukla, R.S., Qin, B. and Cheng, K. (2014) Peptides Used in the Delivery of Small Noncoding RNA. Molecular Pharmaceutics, 11, 3395–3408.

69. Cech, T.R. (2012) The RNA Worlds in Context. CSH Perspectives.

70. Yamazaki, T., Nakagawa, S. and Hirose, T. (2019) Architectural RNAs for Membraneless Nuclear Body Formation. Cold Spring Harb Symp Quant Biol, 84, 227–237.

71. Somasundaram, K., Gupta, B., Jain, N. and Jana, S. (2022) LncRNAs divide and rule: The master regulators of phase separation. Front Genet, 13, 930792.

72. Mattick, J.S., Amaral, P.P., Carninci, P., Carpenter, S., Chang, H.Y., Chen, L.L., Chen, R., Dean, C., Dinger, M.E., Fitzgerald, K.A. et al. (2023) Long non-coding RNAs: definitions, functions, challenges and recommendations. Nat Rev Mol Cell Biol, 24, 430–447.

73. Mattick, J.S. (2023) Enhancers are genes that express organizational RNAs. Front. RNA Res, 1.

74. Schug, K.A. and Lindner, W. (2005) Noncovalent Binding between Guanidinium and Anionic Groups: Focus on Biological-and Synthetic-Based Arginine/Guanidinium Interactions with Phosph[on]ate and Sulf[on]ate Residues. Chemical Reviews, 105, 67–114.

75. Rutledge, L.R., Campbell-Verduyn, L.S. and Wetmore, S.D. (2007) Characterization of the stacking interactions between DNA or RNA nucleobases and the aromatic amino acids. Chemical Physics Letters, 444, 167–175.

76. Brynn Hibbert, D. and Thordarson, P. (2016) The death of the Job plot, transparency, open science and online tools, uncertainty estimation methods and other developments in supramolecular chemistry data analysis. Chemical Communications, 52, 12792–12805.

77. http://supramolecular.org

78. Thordarson, P. (2012) n Supramolecular Chemistry: From Molecules to Nanomaterials Gale, P., Steed, J. (ed.). John Wiley & Sons, Chichester, England, pp. 239–274.

79. Huppert, J.L. and Balasubramanian, S. (2005) Prevalence of quadruplexes in the human genome. Nucleic Acids Research, 33, 2908–2916.

80. Biver, T. (2022) Discriminating between Parallel, Anti-Parallel and Hybrid G-Quadruplexes: Mechanistic Details on Their Binding to Small Molecules. Molecules, 10.3390/molecules27134165.

81. Gu, J., Leszczynski, J. and Bansal, M. (1999) A new insight into the structure and stability of Hoogsteen hydrogen-bonded G-tetrad: an ab initio SCF study. Chemical Physics Letters, 311, 209–214.

82. Wimberger, L., Rizzuto, F.J. and Beves, J.E. (2023) Modulating the Lifetime of DNA Motifs Using Visible Light and Small Molecules. Journal of the American Chemical Society, 145, 2088–2092.

83. Frenkel-Pinter, M., Haynes, J.W., Mohyeldin, A.M., C, M, Sargon, A.B., Petrov, A.S., Krishnamurthy, R., Hud, N.V., Williams, L.D. and Leman, L.J. (2020) Mutually stabilizing interactions between proto-peptides and RNA. Nature Communications, 11, 3137.

84. Xiao, C.-D., Ishizuka, T. and Xu, Y. (2017) Antiparallel RNA G-quadruplex Formed by Human Telomere RNA Containing 8-Bromoguanosine. Scientific Reports, 7, 6695.

85. Mendoza, O., Porrini, M., Salgado, G.F., Gabelica, V. and Mergny, J.L. (2015) Orienting tetramolecular G-quadruplex formation: the quest for the elusive RNA antiparallel quadruplex. Chemistry, 21, 6732–6739.

86. Wang, Y. and Patel, D.J. (1993) Solution structure of a parallel-stranded G-quadruplex DNA. J Mol Biol, 234, 1171–1183.

87. Mathad, R.I., Hatzakis, E., Dai, J. and Yang, D. (2011) c-MYC promoter G-quadruplex formed at the 5′-end of NHE III 1 element: insights into biological relevance and parallel-stranded G-quadruplex stability. Nucleic Acids Research, 39, 9023–9033.

88. Sedoris, K.C., Thomas, S.D., Clarkson, C.R., Muench, D., Islam, A., Singh, R. and Miller, D.M. (2012) Genomic c-Myc quadruplex DNA selectively kills leukemia. Mol Cancer Ther, 11, 66–76.

89. Ghosh, S., Takahashi, S., Ohyama, T., Liu, L. and Sugimoto, N. (2024) Elucidating the Role of Groove Hydration on Stability and Functions of Biased DNA Duplexes in Cell-Like Chemical Environments. Journal of the American Chemical Society.

90. Stump, S., Mou, T.C., Sprang, S.R., Natale, N.A.-O. and Beall, H.A.-O. Crystal structure of the major quadruplex formed in the promoter region of the human c-MYC oncogene.

91. Islam, M.A., Thomas Sd Fau - Murty, V.V., Murty Vv Fau - Sedoris, K.J., Sedoris Kj Fau - Miller, D.M. and Miller, D.M. c-Myc quadruplex-forming sequence Pu-27 induces extensive damage in both telomeric and nontelomeric regions of DNA.

92. Yu, H., Qi, Y., Yang, B., Yang, X. and Ding, Y. (2023) G4Atlas: a comprehensive transcriptome-wide G-quadruplex database. Nucleic Acids Research, 51, D126–D134.

93. Song, J., Gooding, A.R., Hemphill, W.O., Love, B.D., Robertson, A., Yao, L., Zon, L.I., North, T.E., Kasinath, V. and Cech, T.R. (2023) Structural basis for inactivation of PRC2 by G-quadruplex RNA. Science, 381, 1331–1337.

94. Peterson, R.D., Theimer, C.A., Wu, H. and Feigon, J. (2004) New applications of 2D filtered/edited NOESY for assignment and structure elucidation of RNA and RNA-protein complexes. Journal of Biomolecular NMR, 28, 59–67.

95. Iwahara, J., Wojciak Jm Fau - Clubb, R.T. and Clubb, R.T. Improved NMR spectra of a protein-DNA complex through rational mutagenesis and the application of a sensitivity optimized isotope-filtered NOESY experiment.

96. Abramson, J., Adler, J., Dunger, J., Evans, R., Green, T., Pritzel, A., Ronneberger, O., Willmore, L., Ballard, A.J., Bambrick, J. et al. (2024) Accurate structure prediction of biomolecular interactions with AlphaFold 3. Nature, 630, 493–500.

97. He, J., Zhu, S., Liang, X., Zhang, Q., Luo, X., Liu, C. and Song, L.A.-O. LncRNA as a multifunctional regulator in cancer multi-drug resistance.

98. Abdelmohsen, K. and Gorospe, M. (2012) RNA-binding protein nucleolin in disease. RNA Biology, 9, 799–808.

99. Qian, Z., Zhurkin, V.B. and Adhya, S. (2017) DNA–RNA interactions are critical for chromosome condensation in Escherichia coli. Proceedings of the National Academy of Sciences, 114, 12225–12230.

100. Han, F.X., Wheelhouse, R.T. and Hurley, L.H. (1999) Interactions of TMPyP4 and TMPyP2 with Quadruplex DNA. Structural Basis for the Differential Effects on Telomerase Inhibition. Journal of the American Chemical Society, 121, 3561–3570.

101. Burger, A.M., Dai F Fau - Schultes, C.M., Schultes Cm Fau - Reszka, A.P., Reszka Ap Fau - Moore, M.J., Moore Mj Fau - Double, J.A., Double Ja Fau - Neidle, S. and Neidle, S. The G-quadruplex-interactive molecule BRACO-19 inhibits tumor growth, consistent with telomere targeting and interference with telomerase function.

102. Molnár, O.R., Végh, A., Somkuti, J. and Smeller, L. (2021) Characterization of a G-quadruplex from hepatitis B virus and its stabilization by binding TMPyP4, BRACO19 and PhenDC3. Scientific Reports, 11, 23243.

103. Zou, M., Li, J.-Y., Zhang, M.-J., Li, J.-H., Huang, J.-T., You, P.-D., Liu, S.-W. and Zhou, C.-Q. (2021) G-quadruplex binder pyridostatin as an effective multi-target ZIKV inhibitor. International Journal of Biological Macromolecules, 190, 178–188.

104. Fernández, S., Eritja, R., Aviñó, A., Jaumot, J. and Gargallo, R. (2011) Influence of pH, temperature and the cationic porphyrin TMPyP4 on the stability of the i-motif formed by the 5 ′-(C3TA2)4-3 ′ sequence of the human telomere. International Journal of Biological Macromolecules, 49, 729–736.

105. Christou, G.A., Katsiki, N., Blundell, J., Fruhbeck, G. and Kiortsis, D.A.-O. Semaglutide as a promising antiobesity drug.

106. Singleton, M.A.-O. and Eisen, M.B. Evolutionary analyses of intrinsically disordered regions reveal widespread signals of conservation.

107. Levengood, J.D. and Tolbert, B.S. (2019) Idiosyncrasies of hnRNP A1-RNA recognition: Can binding mode influence function. Seminars in Cell & Developmental Biology, 86, 150–161.

108. Portz, B., Lee, B.L. and Shorter, J. (2021) FUS and TDP-43 Phases in Health and Disease. Trends in Biochemical Sciences, 46, 550–563.

109. Yagi, R., Miyazaki, T. and Oyoshi, T. (2018) G-quadruplex binding ability of TLS/FUS depends on the β-spiral structure of the RGG domain. Nucleic Acids Research, 46, 5894–5901.

110. Saric, T., Graef, C.I. and Goldberg, A.L. (2004) Pathway for Degradation of Peptides Generated by Proteasomes: A KEY ROLE FOR THIMET OLIGOPEPTIDASE AND OTHER METALLOPEPTIDASES*. Journal of Biological Chemistry, 279, 46723–46732.

111. Bruno, B.J., Miller Gd Fau - Lim, C.S. and Lim, C.S. Basics and recent advances in peptide and protein drug delivery.

112. Guccione, E. and Richard, S. (2019) The regulation, functions and clinical relevance of arginine methylation. Nature Reviews Molecular Cell Biology, 20, 642–657.

