## Supplementary Information for "Specificity of the stabilising interaction between intrinsically disordered protein sequences and G-quadruplexes in RNA"

### 1. Materials & methods

#### 1.1 General material / instrument use

All solvents and reagents were purchased from chemical supply companies and used without further purification, unless otherwise stated. TERRA G-quadruplex oligonucleotides were synthesised by Dr. Hsiu Lin Li at the UNSW RNA Institute on an OligoPilot 10 DNA/RNA Synthesiser, purified by Strong Anion Exchange High Performance Liquid Chromatography (SAX-HPLC), and desalted using Amicon (R) Ultra Centrifugal Filters. Peptides were synthesised by Solid Phase Peptide Synthesis (SPPS) and purified by Reverse-Phase High Performance Liquid Chromatography (RP-HPLC). Samples were Characterised and purity confirmed by Liquid Chromatography Mass Spectrometry (LC-MS), Nuclear Magnetic Resonance (NMR), Ultraviolet-Visible Spectroscopy (UV-Vis), and Infrared Spectroscopy (IR). Samples used for the TERRA mutant screens were purchased from Integrated DNA Technologies and used without further purification.

#### 1.2 Buffer and other conditions

NMR binding studies were performed in 50 mM sodium/potassium phosphate buffers, with added chloride salts (50 mM) unless otherwise stated, pH adjusted with HCl / NaOH / KOH. Circular dichroism stability studies were performed in Tris Acetate (40 mM Tris, 20 mM acetic acid) pH adjusted to 7.01 with additional acetic acid. All static measurements were obtained at 25 °C, unless otherwise stated.

#### 1.3 NMR Titrations

*NMR titration procedure*: A series of 11 samples were prepared, with a constant concentration of peptide, and increasing equivalents of oligonucleotide. Unless otherwise stated, the peptide concentration was 125 μM, with oligonucleotide concentrations of 0, 12.5, 25, 50, 75, 100, 125, 150, 200, 250 and 500 μM, for a final saturation point of 4x equivalents. Samples were prepared in 160 μL 3 mm NMR tubes in 90% phosphate buffer, 10% $D_2O$. A minimal concentration of 3-(trimethylsilyl)propinoic-2,2,3,3-$d^4$ acid sodium salt was included as an internal standard. Samples were mixed and allowed to sit at room temperature for at least 3 hours before spectra were collected, to allow for folding equilibrium.

*Data analysis*:

The online tool supramolecular.org (1) developed by us (2) used for data analysis. Below is a description of how the data analysis was carried out and the underlying theory and equations used in the supramolecular.org online tool under the “Bindfit” algorithm. Full source code for Bindfit is also available at:

https://github.com/echus/supramolecular-apps

The chemical shift data from the NMR titration was imported into supramolecular.org. A global fit (which greatly improves the fit) model of 3 shifts within the phenylalanine aromatic region was used to produce binding isotherms, as this region was well-defined and exhibited a significant shift. All titrations were run in duplicate, with the weighted average reported. The data was fitted to 1:1 and 2:1 equilibria, with five different binding models considered in each case as detailed below. Preliminary attempts to fit the data to 1:2 binding model did not yield any meaningfully good fit or physically impossible results (e.g. chemical shifts > 100's of ppm) and hence the 1:2 binding model was not considered further in this work. The focus was therefore on comparing the different 2:1 binding model and see if any or all of them where superior to the simple 1:1 model. Overall, the best fitting model in nearly cases was a 2:1 peptide:oligonucleotide statistical fit which is the focus of discussion in the main text. All fitting parameters and models are reported in Supplementary Tables **2-4**. Full details on the equations and terminology used here for the binding models used have been published previously.[3] Below the most important equations referred to in this work here are summarized:

*a) Key terminology:* In the equations below, H = host (peptide), G = guest (RNA or DNA), $[X]_0$ total concentration of species X, and $K$ = association constants) as discussed further in below. N.b. the peptide is defined as the "host" because we are adding RNA or DNA to the peptide solution.

*b) Free energy (ΔG) changes:* The association constants, also be known as the equilibrium constants ($K_a$), for the simple 1:1 H:G complexation as according to equation (S1) can be expressed in free energy ($\Delta G_a$) according to equation (S2) (1).

$$K_\mathrm{a} = \frac{[\mathrm{HG}]}{[\mathrm{H}][\mathrm{G}]} \quad \text{Eq. (S1)} \qquad\qquad \Delta G_\mathrm{a} = -R\mathrm{T}\ln K_\mathrm{a} \quad \text{Eq. (S2)}$$

For 2:1 $H_2G$ complexation is described according to equation (S3) and (S4). We can now express the stepwise association constants ($K_1$ and $K_2$) can be expressed in terms of the free energy changes ($\Delta G_1$ and $\Delta G_1$) according to equations (S5) and (S6) using the microscopic stepwise association constants ($K_{1m}$ and $K_{2m}$) which are derived from the stepwise association constants after correcting for statistical factors as $K_{1m} = K_1/2$ and $K_{2m} = 2K_2$ (4).

$$K_1 = \frac{[\mathrm{HG}]}{[\mathrm{H}][\mathrm{G}]} \quad \text{Eq. (S3)} \qquad\qquad K_2 = \frac{[\mathrm{H_2G}]}{[\mathrm{HG}][\mathrm{H}]} \quad \text{Eq. (S4)}$$

$$\Delta G_1 = -R\mathrm{T}\ln K_{1\mathrm{m}} = -R\mathrm{T}\ln\left(\frac{K_1}{2}\right) \quad \text{Eq. (S5)}$$

$$\Delta G_2 = -R\mathrm{T}\ln K_{2\mathrm{m}} = -R\mathrm{T}\ln(2K_2) \quad \text{Eq. (S6)}$$

*c) Binding models:*

In the data below five different binding models are usually compared.

1. The first on is classical **1:1** equilibria. Here, we define the NMR resonance for the host as $\delta_{\mathrm{H}}$, the guest as $\delta_{\mathrm{G}}$ and the host-guest complex as $\delta_{\mathrm{HG}}$. From this, we can also define the change in resonance for the host-guest complexation as $\delta_{\Delta\mathrm{HG}} = \delta_{\mathrm{HG}} - \delta_{\mathrm{H}}$. If we then define $\delta_0$ = NMR resonance of the host before the guest is added (before the start of titration) we can define the change in in resonance as $\Delta\delta = \delta - \delta_0$. We can now write the NMR version of our simple 1:1 equilibria according to equation (S9) which is derived from the generic quadratic equation used to calculate the concentration of host-guest complex [HG] as previously described (3).

$$\Delta\delta = \frac{\delta_{\Delta\mathrm{HG}}}{[\mathrm{H}]_0}\left(\frac{1}{2}\left\{\left([\mathrm{G}]_0 + [\mathrm{H}]_0 + \frac{1}{K_a}\right) - \sqrt{\left([\mathrm{G}]_0 + [\mathrm{H}]_0 + \frac{1}{K_a}\right)^2 - 4[\mathrm{H}]_0[\mathrm{G}]_0}\right\}\right) \quad \text{Eq. (S7)}$$

2. The second one is the stepwise (non-degenerate) "**full 2:1**" binding model. This model assumes two non-identical two binding sites per molecule of "guest" (RNA) that allows for cooperativity (negative or positive). As with the 1:1 equilibria we first define $\delta_{\Delta\mathrm{HG}_2}$ as the difference between in NMR resonance on the peptide "host" between the 2:1 host-guest complex ($\delta_{\mathrm{H_2G}}$) and the host NMR resonance ($\delta_{\mathrm{H}}$), that is $\delta_{\Delta\mathrm{H_2G}} = \delta_{\mathrm{H_2G}} - \delta_{\mathrm{H}}$. Using $\delta_{\Delta\mathrm{HG}} = \delta_{\mathrm{HG}} - \delta_{\mathrm{H}}$ for the change in NMR resonance for the 1:1 complex formation and the observed change in resonance as $\Delta\delta = \delta - \delta_0$ as before with the 1:1 equilibria, we obtain equation (S8).

$$\Delta\delta = \frac{\delta_{\Delta\mathrm{HG}}K_1[\mathrm{G}]_0[\mathrm{H}] + 2\delta_{\Delta\mathrm{H_2G}}K_1K_2[\mathrm{G}]_0[\mathrm{H}]^2}{[\mathrm{H}]_0(1 + K_1[\mathrm{H}] + K_1K_2[\mathrm{H}]^2)} \quad \text{Eq. (S8)}$$

Here the guest [H] concentration is obtained from the cubic equation (S9) (3).

$$[\mathrm{H}]^3(K_1K_2) + [\mathrm{H}]^2\{K_1(2K_2[\mathrm{G}]_0 - K_2[\mathrm{H}]_0 + 1)\} + [\mathrm{G}]\{K_1([\mathrm{G}]_0 - [\mathrm{H}]_0) + 1\} - [\mathrm{H}]_0 = 0 \quad \text{Eq. (S9)}$$

Notably, for the "**full 2:1**" model we make no assumptions about the correlation between either $K_1$ and $K_2$ ($K_1 \neq 4K_2$) or $\delta_{\Delta\mathrm{H_2G}}$ and $\delta_{\Delta\mathrm{HG}}$ ($\delta_{\Delta\mathrm{H_2G}} \neq 2\delta_{\Delta\mathrm{HG}}$).

3. The third one is the stepwise (non-degenerate) "**additive 2:1**" binding model. To reduce the number of fitted parameters we note that in many circumstances it can be assumed that the induced chemical shifts of the protons being monitored in the NMR experiment are simply additive, *i.e.*, for proton resonance Y, the shift caused by the second binding event is exactly the same as from the first binding. It then follows that $\delta_{\Delta H_2G} = 2\delta_{\Delta HG}$ and we can simplify equation (S8) to yield equation (S10).

$$\Delta\delta = \frac{\delta_{\Delta HG} K_1 [G]_0 [H]\{1+2K_2[H]\}}{[H]_0(1+K_1[H]+K_1K_2[H]^2)} \quad \text{Eq. (S10)}$$

We have for the "**additive 2:1**" model therefore made the assumption that $\delta_{\Delta H_2G} = 2\delta_{\Delta HG}$, whilst not making any assumptions about the correlation between either $K_1$ and $K_2$ ($K_1 \neq 4K_2$).

4. The forth model is the stepwise "**non-cooperative 2:1**" model. Here we revert back to noting that the chemical shift differences between the first and second binding may not be correlated ($\delta_{\Delta H_2G} \neq 2\delta_{\Delta HG}$) but instead we make the assumption that the 2:1 binding is non-cooperative and therefore that $K_1 = 4K_2$. We can then use this to rewrite equation (S8) and replace $K_2$ with $K_1/4$ to obtain equation (S11). If desired, $K_2$ can be calculated back from $K_1$ as $K_2 = K_1/4$.

$$\Delta\delta = \frac{\delta_{\Delta HG} K_1 [G]_0 [H]\left\{1+\frac{\delta_{\Delta H_2G} K_1 [H]}{2}\right\}}{[H]_0\left(1+K_1[H]+\frac{(K_1[H])^2}{2}\right)} \quad \text{Eq. (S11)}$$

We have in "**non-cooperative 2:1**" model made the assumption that $K_1 = 4K_2$ whilst making no assumption about the correlation between ($\delta_{\Delta H_2G}$ and $\delta_{\Delta HG}$ (($\delta_{\Delta H_2G} \neq 2\delta_{\Delta HG}$).

5. The fifth model is the "**statistical 2:1**" model. Here we not only make the assumption that the binding is non-cooperative ($K_1 = 4K_2$) but also that the chemical shift changes are simply additive ($\delta_{\Delta H_2G} = 2\delta_{\Delta HG}$. This means in other words we make the assumption that the two binding site behave like two independent hosts. This leads to a further simplification of equation (S8) to equation (S12).

$$\Delta\delta = \frac{\delta_{\Delta HG} K_1 [G]_0 [H]\left\{1+\frac{K_1 [H]}{2}\right\}}{[H]_0\left(1+K_1[H]+\frac{(K_1[H])^2}{2}\right)} \quad \text{Eq. (S12)}$$

In this "**statistical 2:1**" model we have therefore made the assumptions that $K_1 = 4K_2$ and that ($\delta_{\Delta H_2G} = 2\delta_{\Delta HG}$. It should also be noted that in this situation, the data could also be fitted to the simple **1:1** model according to Equation (S8) by simply multiply the total host concentration $[H]_0$ by a factor of 2.

The resulting association constant $K_a$ is then equal to the non-cooperative microscopic binding constants, *i.e.*, $K_a = K_{1m} = K_{2m}$, which means $K_1 = K_a/2$ and $K_2 = 2K_a$. As discussed, the statistical 2:1 model always gave the best fit and hence for brevity in some of the Figures further below, only the $K_1$ values are shown as $K_2$ can easily calculated from with $K_2 = K_1/4$.

d) *Comparing the models:*

To analyze and compare the model used we used the Bayesian Information Criteria (BIC) (5) as a robust method for model comparison. The challenge is that the more complicated the model (= higher number of parameters), the better the fit is likely to be. However, generally we should only pick a more complicated model if the fit is significantly better when compared to a simpler model. The BIC based on the calculated log-likelihood, the number of parameters, and the number of data points, whereby an increase in the number of parameters leads to a penalty (increase) in the BIC value. A low BIC is generally better and when comparing two models, and a difference of more than 6-10 is usually considered as strong evidence that there is a significant difference between the two models being compared.

#### 2. Synthesis

##### 2.1 Synthesis of oligonucleotides

Oligonucleotide synthesis

Synthesis of TERRA DNA/RNA oligonucleotides was performed at the UNSW RNA Institute on a Cytiva ÄKTA Oligopilot™ 10 Plus oligonucleotide synthesiser. Synthesis was performed on preloaded Cytiva primer support Ribo 300 polymer resin in a 1.2 mL stainless steel column. Synthesis was carried out under an atmosphere of dry argon. Phosphoramidites were purchased from ThermoFisher and used without further purification. The solid-phase resin was purchased from Cytiva. The remaining reagents were purchased from Sigma Aldrich and Tedia High Purity Solvents and used without additional purification. SAX-HPLC purification was performed as per section 1.5.1. Millipore Amicon® Ultra-0.5 3kDa centrifugal filters were purchased from Sigma and used for desalting.

Mutant TERRA oligonucleotides were purchased from Integrated DNA technologies and used without further purification. LC-MS was performed to ensure purity as per section 1.5.3.

***Supplementary Table 1:*** G-quadruplex RNA & DNA sequences used

| **Code** | **Sequence (5’-3’)** |
|---|---|
| TERRA ($G_3$UUA) | 5’-GGGUUAGGGUUAGGGUUAGGG-3’ |
| G4 DNA | 5’-GGGTTAGGGTTAGGGTTAGGG-3’ |
| $G_3U_3$ | 5’-GGGUUUGGGUUUGGGUUUGGG-3’ |
| $G_3U_2$ | 5’-GGGUUGGGUUGGGUUGGG-3’ |
| $G_3U_1$ | 5’-GGGUGGGUGGGUGGG-3’ |
| $G_3A_3$ | 5’-GGGAAAGGGAAAGGGAAAGGG-3’ |
| $G_3A_2$ | 5’-GGGAAGGGAAGGGAAGGG-3’ |
| $G_3A_1$ | 5’-GGGAGGGAGGGAGGG-3’ |

#### 2.2 Synthesis of peptides

Solid Phase Peptide Synthesis (SPPS) of **RGGFGGRGG**

SPPS reagents and solvents were purchased from ChemSupply and Sigma Aldrich. Fmoc-protected amino acids were purchased from Chem-Impex. No additional purification was necessary. **RGGFGGRGG** was successfully synthesised by solid phase peptide synthesis (SPPS). 2-chlorotrityl chloride resin (0.400 g, 0.446 mmol) was added to a 12 mL polypropylene syringe, before being washed and swelled with dichloromethane (6 mL, 2 x 1 min, 1 x 15 min). Fmoc-Gly-OH (0.401 g, 1.338 mmol) was dissolved in *N*,*N*-diisopropylethylamine (0.62 mL, 3.57 mmol), peptide-grade *N*,*N*-dimethylformamide (2 mL) and dichloromethane (2 mL). The amino acid solution was added to the syringe and left to load with shaking at room temperature (18 h). Excess reagents were ejected and the resin was washed with dichloromethane (6 mL, 2 x 1 min) and N,N-dimethylformamide (6 mL, 3 x 1 min). The Fmoc protecting group was removed in a deprotecting solution of piperidine in *N*,*N*-dimethylformamide (20%, v/v, 6 mL, 1 x 5 min, 1 x 10 min), followed by washes with N,N-dimethylformamide (6 mL, 5 x 1 min). The remaining eight amino acids, Fmoc-Gly-OH (0.401 g, 1.34 mmol), Fmoc-L-Arg(Pbf)-OH (0.870 g, 1.34 mmol), Fmoc-Gly-OH (0.403, 1.34 mmol), Fmoc-Gly-OH (0.401 g, 1.34 mmol), Fmoc- L-Phe-OH (0.520 g, 1.34 mmol), Fmoc-Gly-OH (0.396 g, 1.34 mmol), Fmoc-Gly-OH (0.395 g, 1.34 mmol) and Fmoc-L-Arg(Pbf)-OH (0.870 g, 1.34 mmol) were stepwise dissolved in N,N-diisopropylethylamine (0.47 mL, 2.68 mmol), and a coupling solution of 1-hydroxybenzotriazole hydrate and *N*,*N*,*N*',*N*'-tetramethyl-*O*-(1*H*-benzotriazole-1-yl) uronium hexafluorophosphate in peptide-grade *N*,*N*-dimethylformamide (2.68 mL, 0.5 M). The amino acid solution was injected into the syringe for each coupling step, and shaken at room temperature (60 min), with a Kaiser test performed between each step to assess the efficiency of coupling. Each coupling was followed by a deprotection in piperidine in *N*,*N*-dimethylformamide (20% v/v, 6 mL, 1 x 5 min, 1 x 10 min) and *N*,*N*-dimethylformamide (6 mL, 5 x 1 min). The final deprotection was followed by washes in *N*,*N*-dimethylformamide (6 mL, 5 x 1 min), dichloromethane (6 mL, 3 x 1 min) and methanol (6 mL, 3 x 1 min), and the resin was dried under nitrogen. The peptide was cleaved from the resin in a solution of trifluoroacetic acid (95% v/v, 5.7 mL), triisopropyl silane (2.5% v/v, 0.15 mL) and Milli-Q™ water (2.5% v/v, 0.15 mL), with shaking at room temperature (3 h). The contents of the syringe were ejected into ice cold diethyl ether (17.5 mL) and centrifuged (4000 rpm, 5 mL). The supernatant was discarded with the resulting precipitate washed and centrifuged twice in additional ice cold diethyl ether (17.5 mL). The precipitate was dissolved in a minimal amount of water and freeze dried, giving the crude product as an off-white powder. The crude product was dissolved in Milli-Q™ water and HPLC-grade acetonitrile (1:1 v/v) and purified by HPLC (Section 1.5.1), affording the final peptide as a white fluffy solid (164.2 mg, 0.200 mmol, 45%). HRMS (ESI$^+$) *m/z*: $[M+H]^+$ calculated for $C_{33}H_{55}N_{15}O_{10}$: 821.4256, found:

821.4247. $^1$H NMR (600 MHz, $D_2O$) δ 7.33 – 7.17 (m, 5H, Phe-Ar), 4.59 – 4.52 (m, 1H, Arg-a-CH), 4.30 (dd, J = 8.9, 5.4 Hz, 1H, Arg-a-CH), 4.04 (t, J = 6.5 Hz, 1H, Phe-a-CH), 4.00 – 3.91 (m, 2H, a-CH), 3.93 – 3.75 (m, 11H Gly-$CH_2$), 3.14 (dt, J = 10.9, 6.9 Hz, 4H, Arg-$CH_2$), 3.10 (dd, J = 13.9, 6.5 Hz, 1H, Arg $CH_2$), 2.97 (dd, J = 13.9, 8.6 Hz, 1H, Phe-b-CH), 1.95 – 1.81 (m, 2H, Arg-b-$CH_2$), 1.78 – 1.68 (m, 1H, Arg g-$CH_2$), 1.68 – 1.51 (m, 3H, Arg d-$CH_2$). HPLC: $t_r$ = 8.5 min (10-95%, $CH_3CN$:$H_2O$ gradient, 35 min, 1.0 mL/min).

#### 3. Peptide characterisation

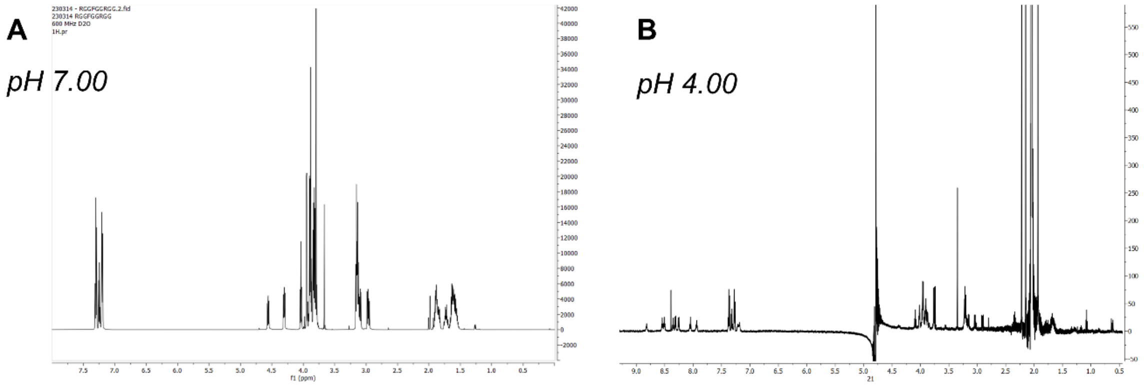

***Supplementary Figure 1:*** *$^{1}H$ NMR spectra (600 MHz, 25 °C, 90% $H_2O$, 10% $D_2O$) of **RGGFGGRGG** peptide (10 mM) at (A): pH 7, (B): pH 4.*

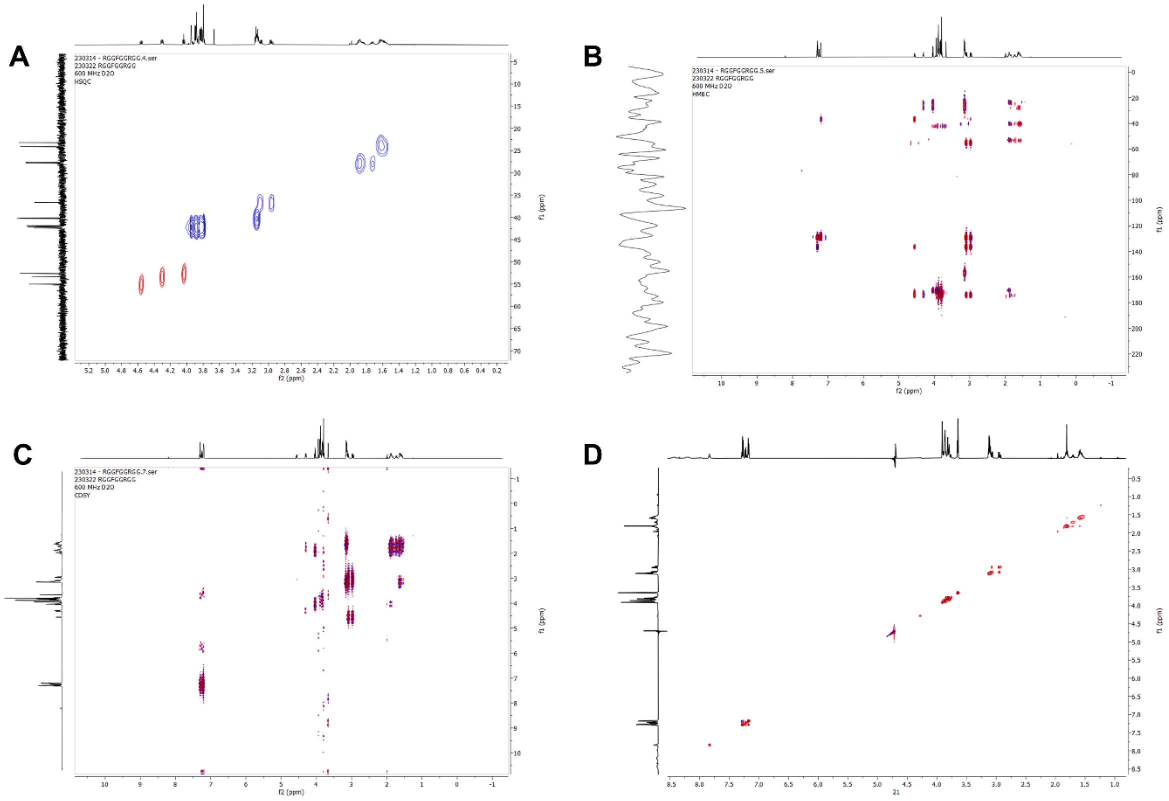

***Supplementary Figure 2:*** *2-dimensional NMR characterisation of the **RGGFGGRGG** peptide at pH 7 (600 MHz, 25 °C, 90% $H_2O$, 10% $D_2O$. 10 mM peptide, 50 mM sodium phosphate buffer (pH 7.01). (A): $^{1}H$-$^{13}C$ HSQC NMR spectra, (B): $^{1}H$-$^{13}C$ HMBC NMR spectra, (C): $^{1}H$-$^{1}H$ HSQC NMR spectra, (D): $^{1}H$-$^{1}H$ NOESY NMR spectra.*

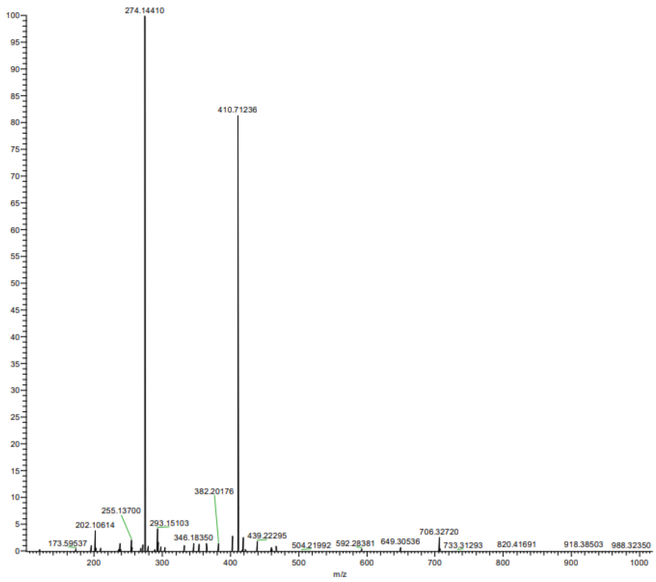

***Supplementary Figure 3:*** *High resolution mass spectrum of* ***RGGFGGRGG*** *peptide, confirming monoisotopic mass.*

### 4. Oligo characterisation

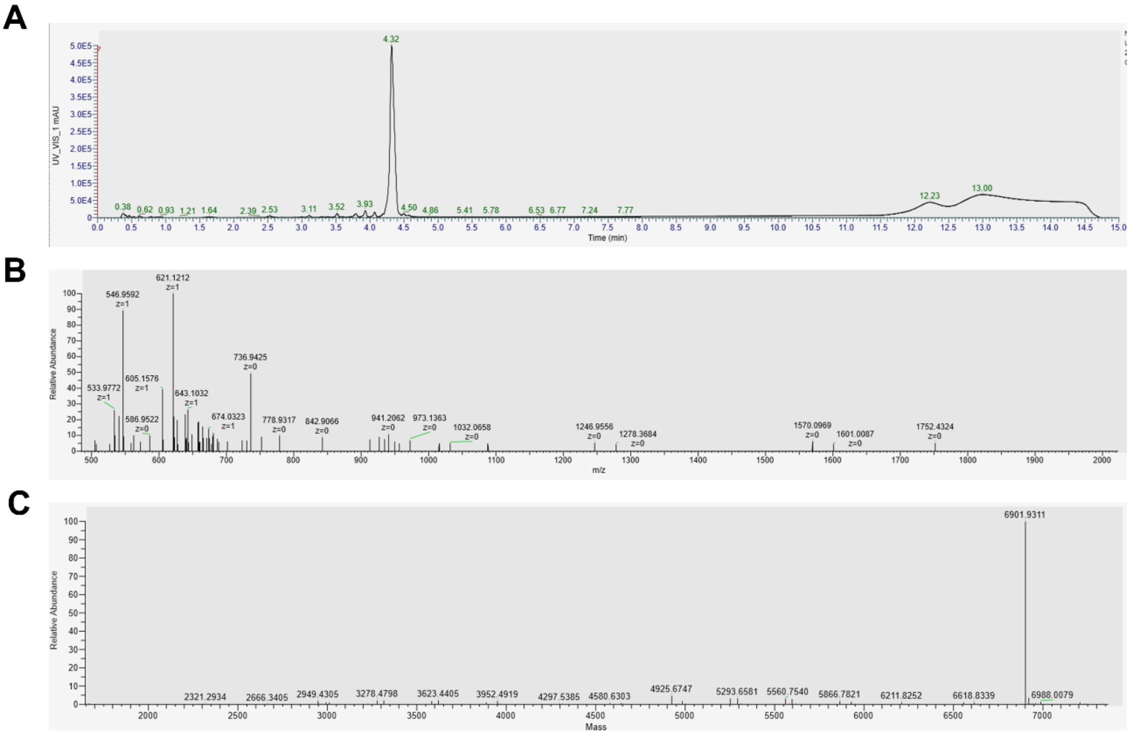

***Supplementary Figure 4:*** *Characterisation of 5’-GGG UUA GGG UUA GGG UUA GGG-3’ (TERRA) RNA. (A): Analytical HPLC spectrum. (B): Mass spectrum trace. (C): Deconvoluted mass spectrum trace.*

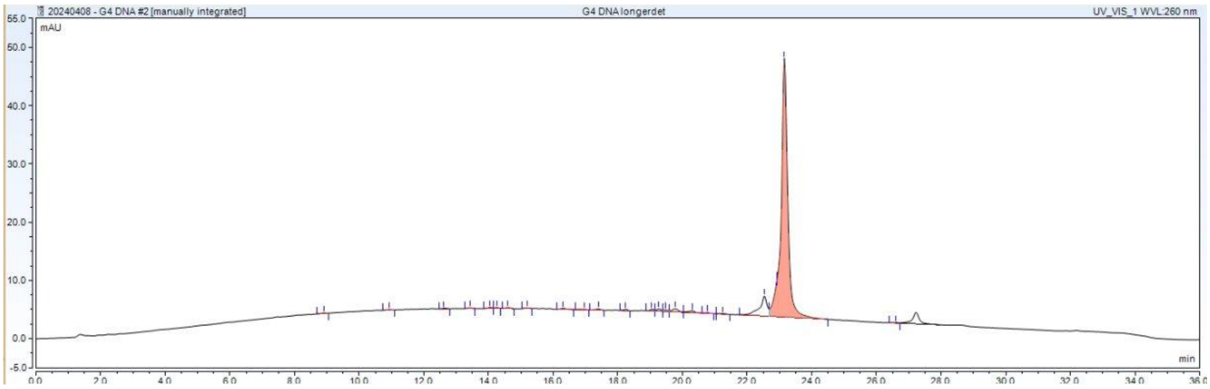

***Supplementary Figure 5:*** *Analytical HPLC spectrum of 5’-GGG TTA GGG TTA GGG TTA GGG-3’ (TERRA) DNA.*

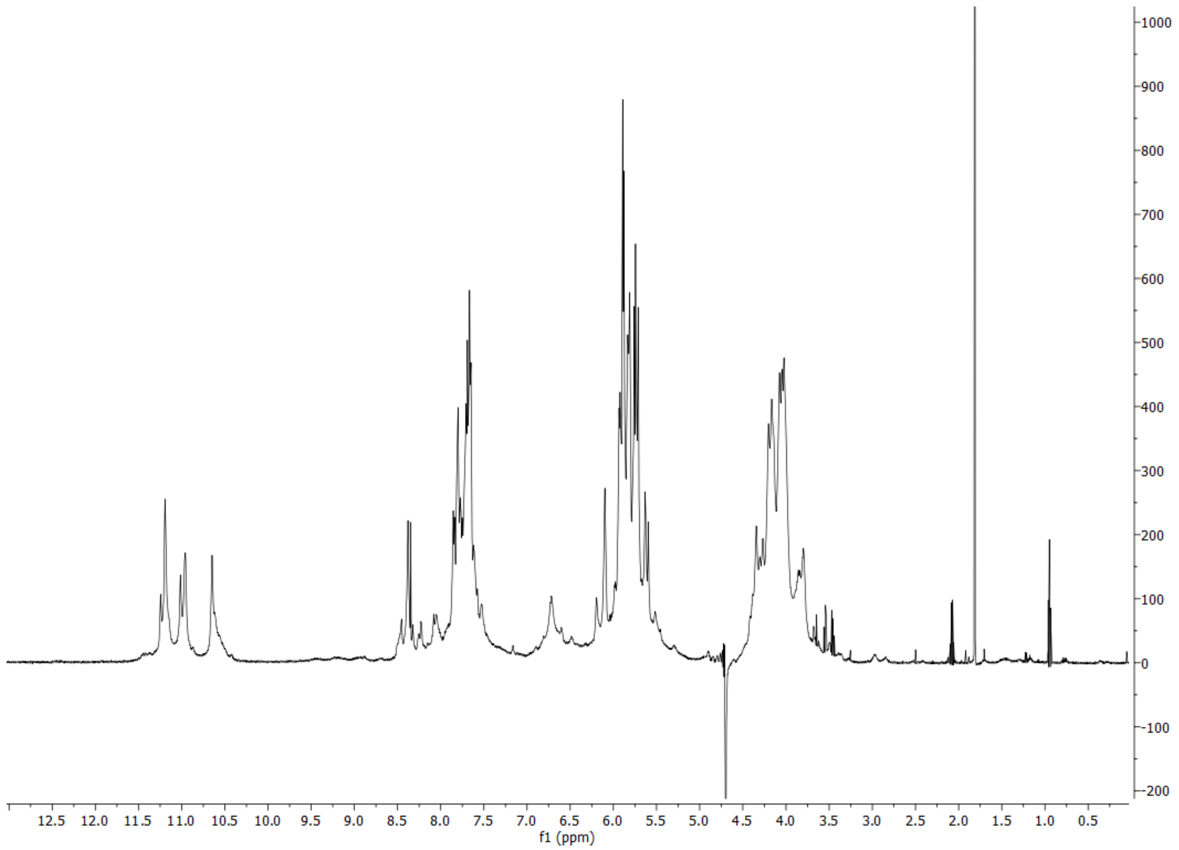

***Supplementary Figure 6:*** *[1]H NMR spectrum (600 MHz, 25 °C, 90% $H_2O$, 10% $D_2O$) of folded 5'-GGG UUA GGG UUA GGG UUA GGG-3' (TERRA) RNA. 1 mM RNA, 50 mM Potassium Phosphate, 50 mM Potassium Chloride (pH 7.01). Imino exchange peaks at 10.5-11.5 ppm confirm folding of a parallel G-quadruplex structure.*

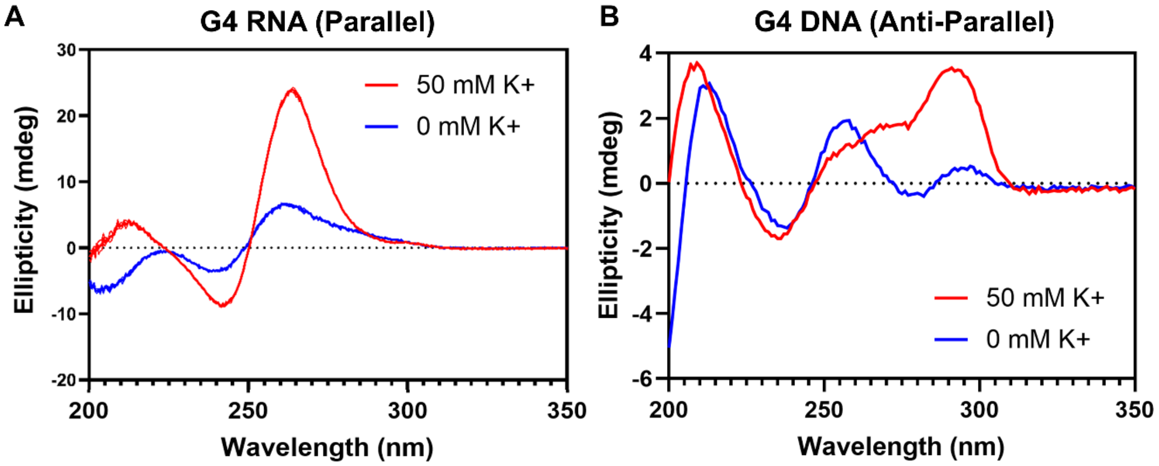

***Supplementary Figure 7:*** *Circular dichroism spectra showing the folding of G-quadruplex structures upon the addition of 50 mM KCl. (A): 3'-GGG UUA GGG UUA GGG UUA GGG-5' (TERRA) RNA folds into a parallel G4 structure (positive band at 260 nm). (B): 3'-GGG TTA GGG TTA GGG TTA GGG-5' DNA folds into an antiparallel G4 structure (positive band at 294 nm).*

#### 5. Supplemental binding data

Raw NMR titration data can be found on GitHub.

***Supplementary Table 2:*** *Association constants of folded and unfolded TERRA G-quadruplex RNA and DNA, derived from NMR titrations, fit to a range of 1:1 and 2:1 (peptide:oligonucleotide) fitting models as explained in Section 1.9 above. $K_1$ = association constant for first binding event ($M^{-1}$). $K_2$ = association constant for second binding event. BIC = Bayesian Information Criterion, a statistic used for fitting model selection. Lower (more negative) numbers are indicative of a more successful fitting model. Shown in bold is the statistical 2:1 model, which is the best fitting model for the majority of titrations and for consistency, all the data in our work is then compared using this model.*

| | ***RNA*** | | | ***DNA*** | | |
|---|---|---|---|---|---|---|
| *Binding Model* | $K_1$ | $K_2$ | BIC | $K_1$ | $K_2$ | BIC |
| **G-quadruplex (folded)** | | | | | | |
| *1:1* | 6.76 x10$^5$ | n/a | -110 | 9.12 x10$^2$ | n/a | -352 |
| *Full 2:1* | 1.47 x10$^1$ | 7.52 x10$^6$ | -147 | 1.07 x10$^0$ | 5.59 x10$^6$ | -283 |
| *Additive 2:1* | 1.55 x10$^4$ | 3.67 x10$^4$ | -178 | 3.41 x10$^1$ | 4.62 x10$^4$ | -348 |
| *Non-Coop 2:1* | 2.20 x10$^4$ | 5.50 x10$^3$ | -173 | 7.81 x10$^2$ | 1.95 x10$^2$ | -342 |
| *Statistical 2:1* | **4.46 x10$^4$** | **1.12 x10$^4$** | **-236** | **7.86 x10$^2$** | **1.97 x10$^2$** | **-352** |
| **G-quadruplex (unfolded)** | | | | | | |
| *1:1* | 9.51 x10$^6$ | n/a | -80 | 4.46 x10$^4$ | n/a | -193 |
| *Full 2:1* | 1.61x10$^1$ | 4.56x10$^6$ | -99 | 2.11 x10$^2$ | 3.03 x10$^7$ | -156 |
| *Additive 2:1* | 1.45 x10$^5$ | 2.64 x10$^4$ | -123 | 2.48 x10$^4$ | 5.16 x10$^2$ | -193 |
| *Non-Coop 2:1* | 1.82 x10$^4$ | 4.56 x10$^3$ | -130 | 1.91 x10$^3$ | 4.78 x10$^2$ | -189 |
| *Statistical 2:1* | **1.23 x10$^4$** | **3.08 x10$^3$** | **-70** | **7.90 x10$^3$** | **1.97 x10$^3$** | **-194** |

***Supplementary Table 3:*** *Association constants derived from NMR titrations, fit to a 2:1 (peptide:oligonucleotide) statistical fitting model, including free energy of the interaction.*

| | ***RNA*** | | | ***DNA*** | | |
|---|---|---|---|---|---|---|
| *Oligonucleotide Species* | $K_1$ | $K_2$ | Free Energy (kJ/mol) | $K_1$ | $K_2$ | Free Energy (kJ/mol) |
| ***G-quadruplex (Folded)*** | 4.5 x10$^4$ | 1.1 x10$^3$ | -24.8 | 7.9 x10$^2$ | 2.0 x10$^2$ | -14.8 |
| ***G-quadruplex (Unfolded)*** | 1.2 x10$^4$ | 3.1 x10$^3$ | -21.6 | 7.9 x10$^3$ | 2.0 x10$^3$ | -20.5 |

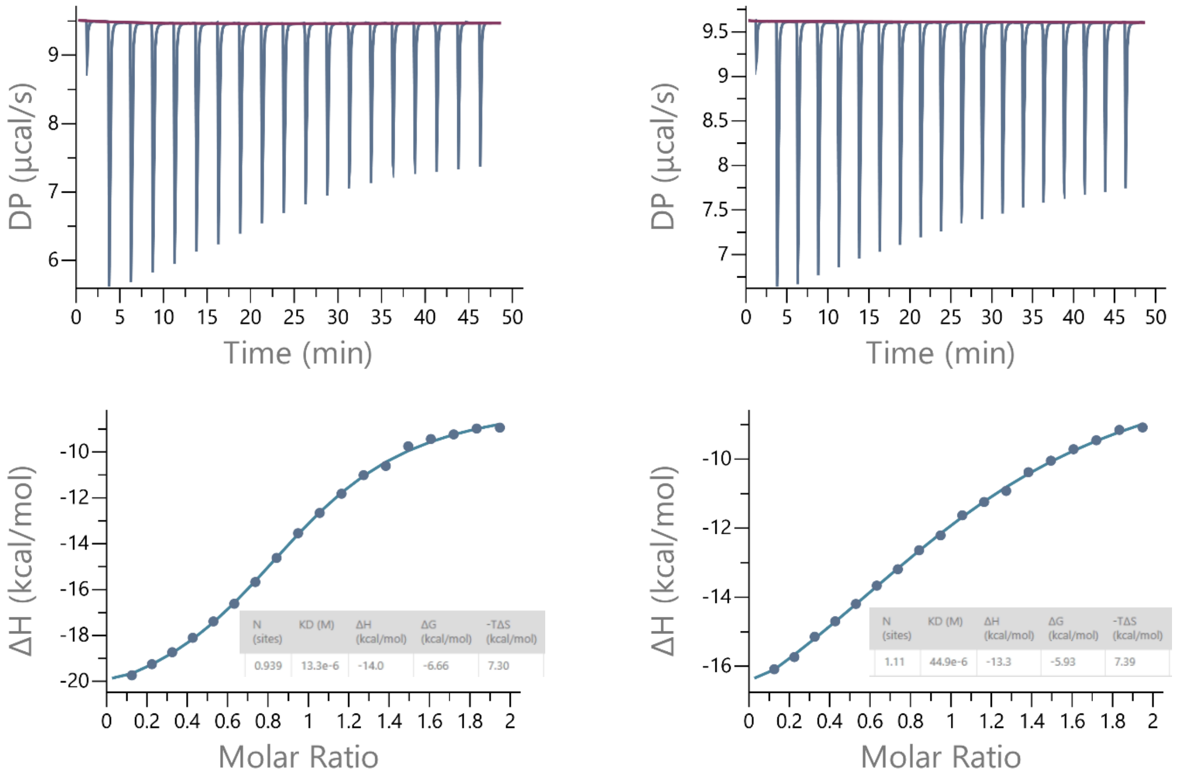

***Supplementary Figure 8:*** *Isothermal Titration Calorimetry (ITC**)** binding isotherm of 5'-GGG UUA GGG UUA GGG UUA GGG-3' (TERRA) RNA with the **RGGFGGRGG** peptide, in duplicate. Average:* $K_1$ *= 48730* $M^{-1}$*. Association constant is in good correlation with NMR titration data.*

#### 6. G-quadruplex folding studies

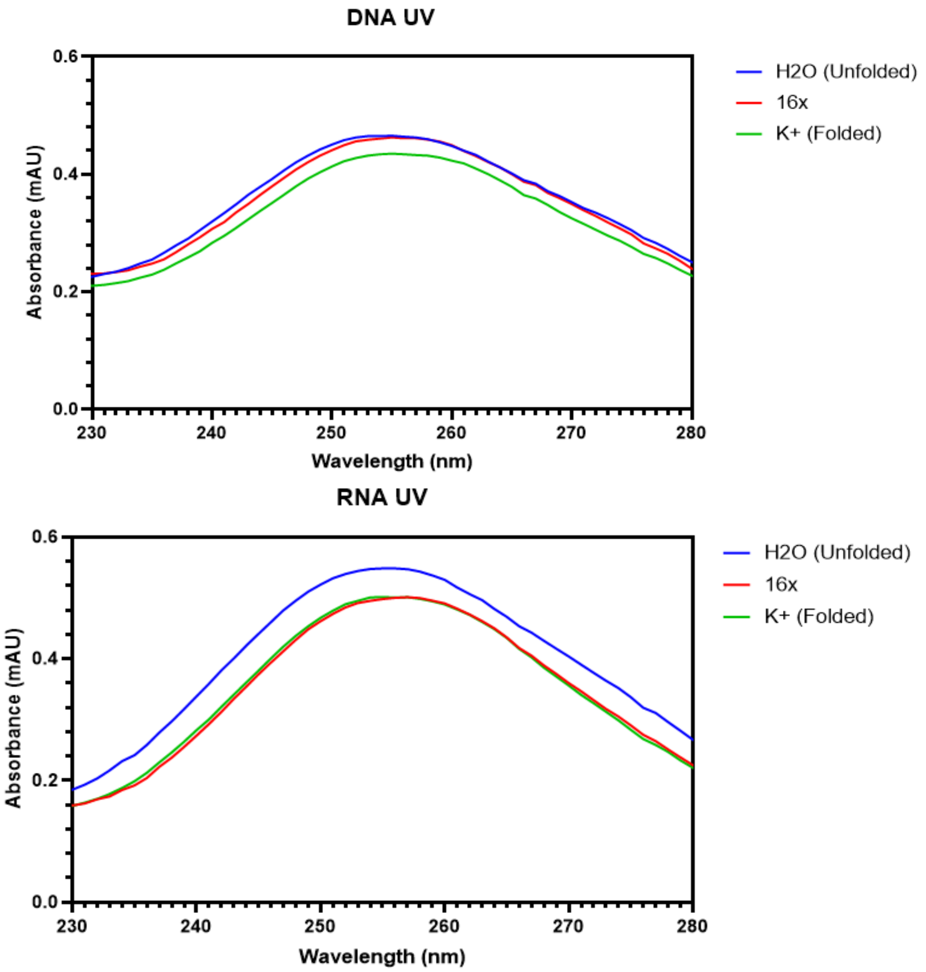

***Supplementary Figure 9:*** *Ultraviolet-Visible spectra of (top): 5'-(GGGUUA)$_3$GGG-3' (TERRA G4) RNA & (bottom): 5'-(GGGTTA)$_3$GGG-3' (TERRA G4) DNA. Blue line shows single-stranded control oligonucleotide sequence. Red line shows shift upon addition of* ***RGGFGGRGG*** *peptide. Green line shows shift upon addition of KCl. In both RNA & DNA, the spectra undergoes a bathochromic and hypochromic shift upon addition of 16 equivalents KCl, indicating G-quadruplex formation. Additions of 16 equivalents of the* ***RGGFGGRGG*** *peptide only successfully fold the RNA, leaving DNA unfolded.*

#### 7. RGGFGGRGG with parallel & antiparallel DNA G4s

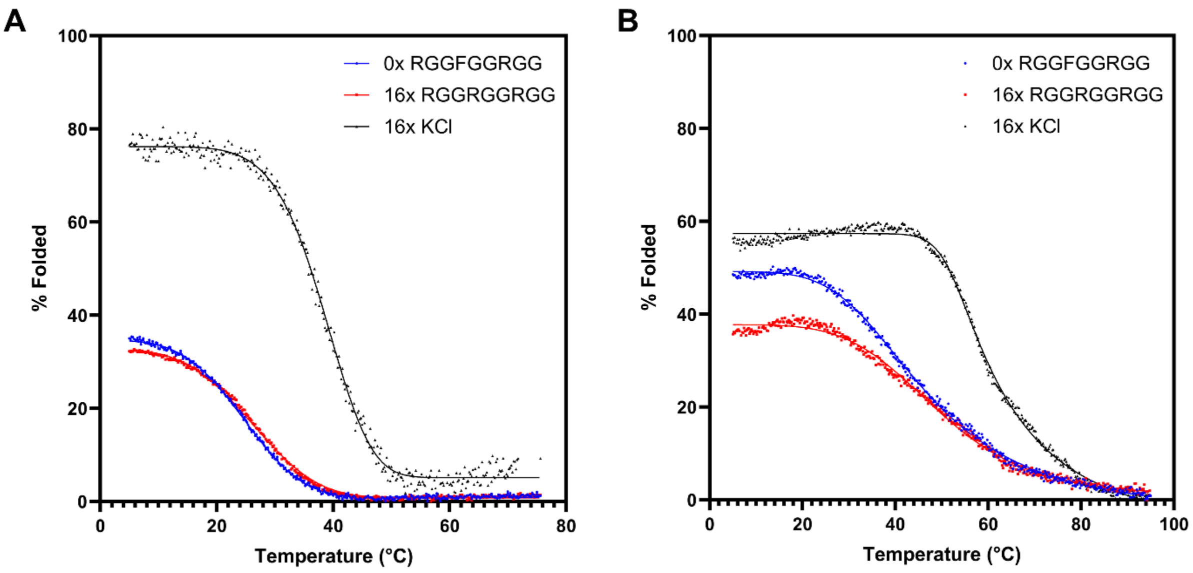

***Supplementary Figure 10:*** *Circular dichroism melting curve of upon additions of* ***RGGFGGRGG*** *peptide. 20 μM RNA, 1x Tris Acetate buffer @ pH 7.0. Data is baseline corrected, normalised for concentration by a UV reading at A260, and scaled to a 100% folded control (oligo in the presence of 100 mM KCl). In both cases, the* ***RGGFGGRGG*** *peptide does not stabilise or template the G4 structure.* ***a:*** *5'-GGG TTA GGG TTA GGG TTA GGG-3' (Antiparallel) DNA G4.* ***b:*** *5'-GGG AA GGG AA GGG AA GGG-3' (Parallel) DNA G4.*

#### 8. TERRA mutant screens

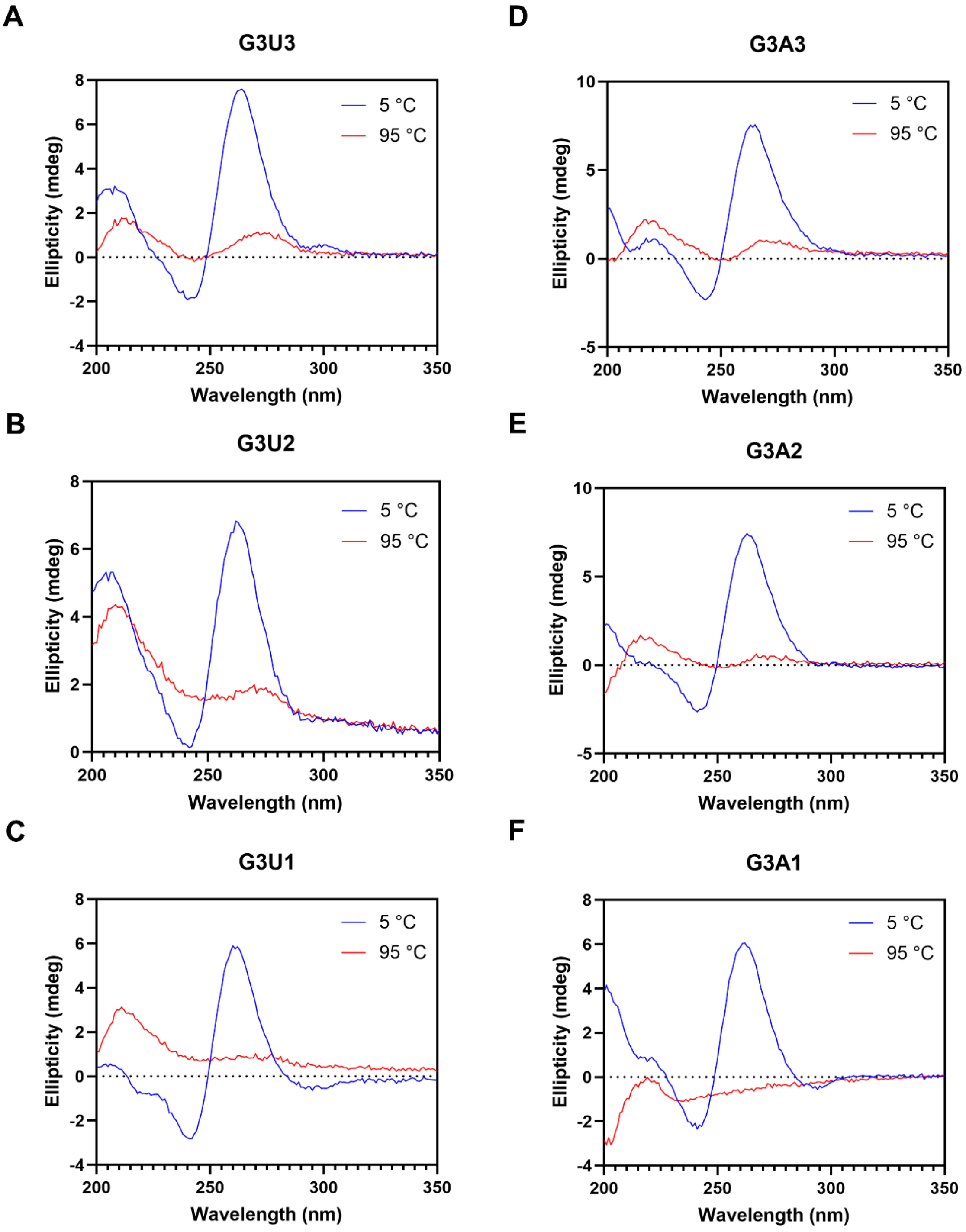

***Supplementary Figure 11:*** *Circular Dichroism spectra of mutated G-quadruplex forming sequences, at 5 °C (blue), and 95 °C (red), in the presence of KCl (100 mM), Potassium Phosphate (pH 7.01, 50 mM). All tested sequences are forming parallel G-quadruplex structures, by the intense positive band at 260 nm.*

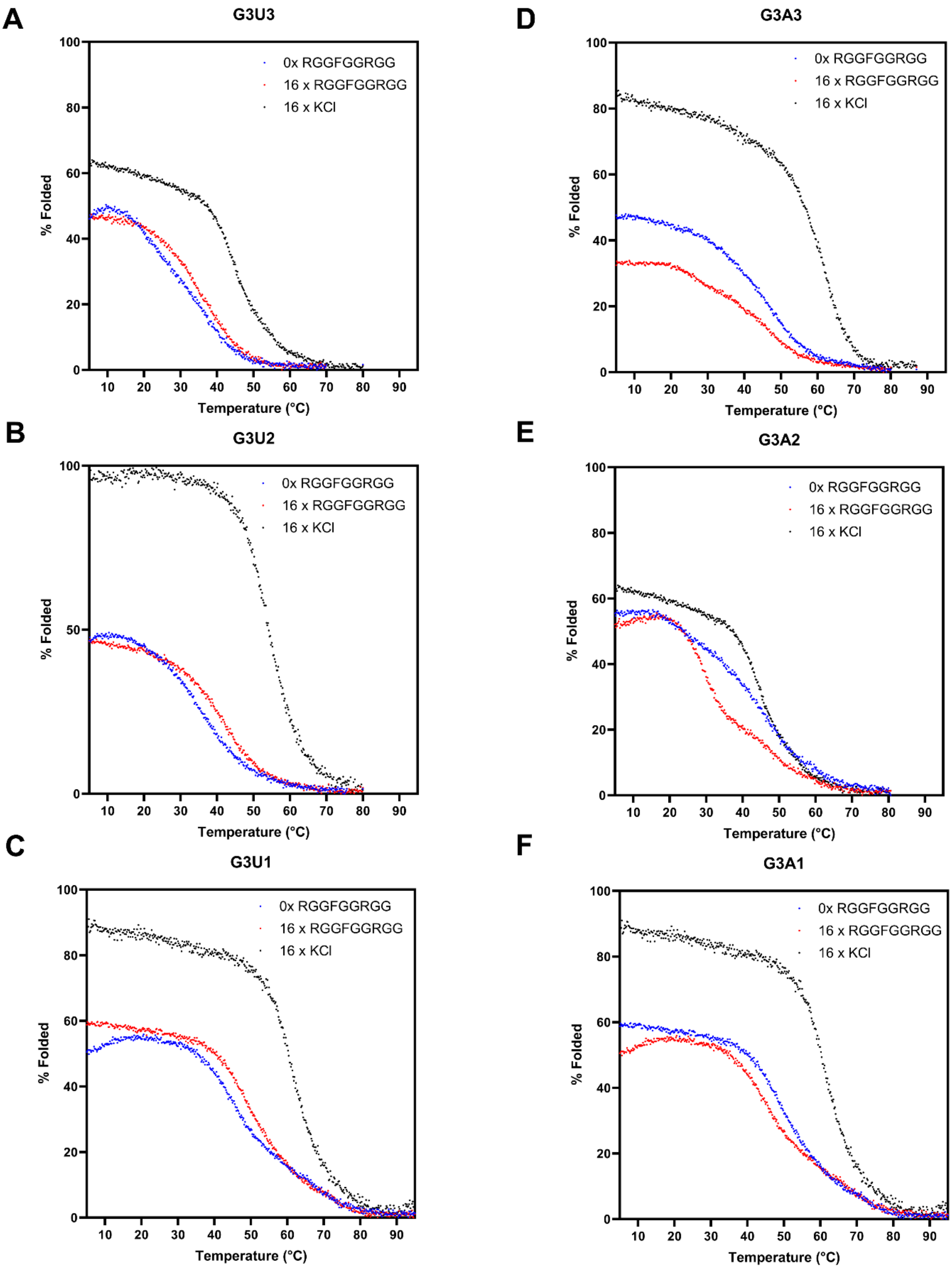

***Supplementary Figure 12:*** *Circular Dichroism melting point experiments of mutated G-quadruplex forming sequences, in the absence of folding salts (blue), 16 equivalents of the* ***RGGFGGRGG*** *peptide (red), and 16 equivalents of the endogenous ligand, potassium (black). 20 µM RNA, 1x Tris Acetate buffer @ pH 7.0. Data is baseline corrected, normalised for concentration by a UV reading at A260, and scaled to a 100% folded control (oligo in the presence of 100 mM KCl). The* ***RGGFGGRGG*** *peptide very minorly stabilizes* $G_3U_{1-3}$ *sequences, but has no stabilizing effect on* $G_3A_{1-3}$*. None of the mutants are templated by* ***RGGFGGRGG****.*

#### 9. Negative control peptides

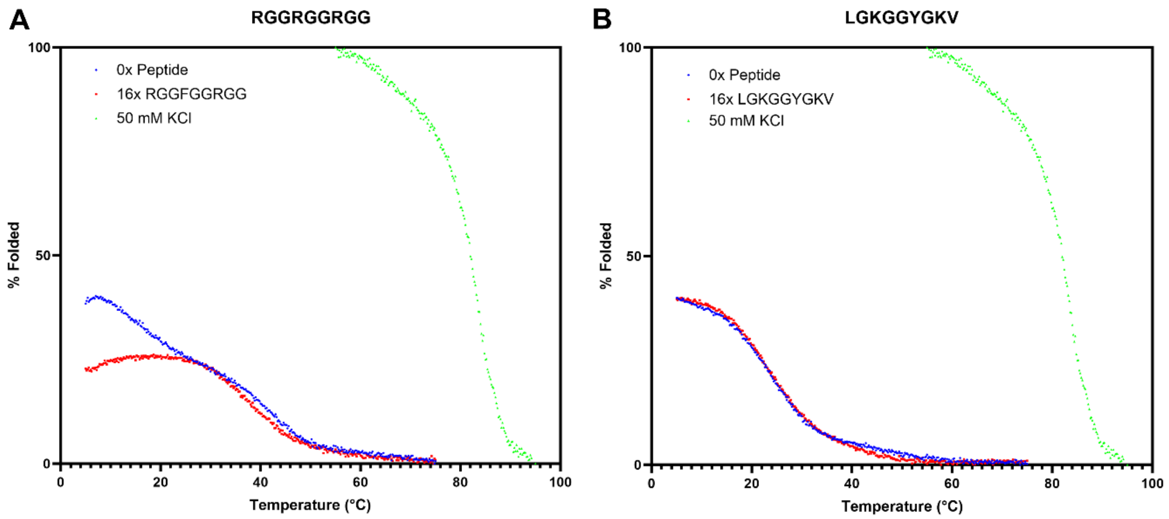

***Supplementary Figure 13:*** *Circular Dichroism melting point experiments of the TERRA G-quadruplex sequence 5'-GGG UUA GGG UUA GGG UUA GGG-3', in the absence of folding salts (blue), 16 equivalents of peptide (red), and 50 mM KCl (green). 20 µM RNA, 1x Tris Acetate buffer @ pH 7.0. Data is baseline corrected, normalised for concentration by a UV reading at A260, and scaled to a 100% folded control (oligo in the presence of 100 mM KCl). (A): The mutated peptide without phenylalanine **RGGRGGRGG** destabilizes the TERRA G-quadruplex structure. (B): A non-interacting peptide, **LGKGGYGKV** has no impact on the G-quadruplex structure. Raw NMR titration data of **LGKGGYGKV** with TERRA RNA available on GitHub.*

#### 10.Structural NMR spectra

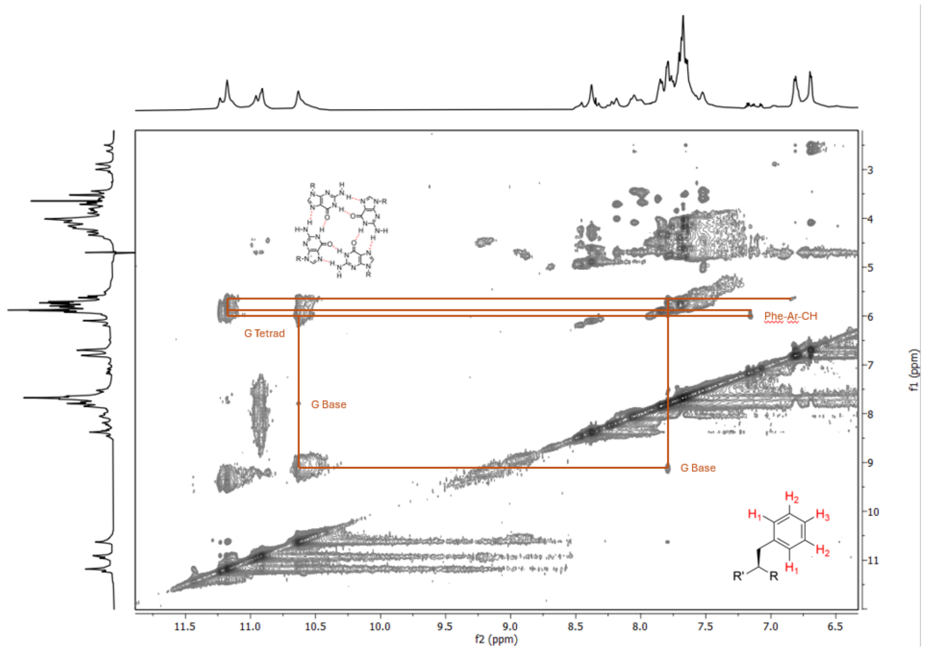

***Supplementary Figure 14:*** *[1]H-[1]H NOESY NMR spectra (600 MHz, 25 °C, 90% $H_2O$, 10% $D_2O$) of* ***RGGFGGRGG*** *peptide and 5'-GGG UUA GGG UUA GGG UUA-3' (TERRA) RNA at pH 7. 2 mM peptide, 2 mM RNA 50 mM Sodium Phosphate (pH 7.01). Interactions: Arginine and RNA Uracil/Guanine Sugars. Phenylalanine and RNA Guanine Sugars. Interactions: Peptide aromatic phenylalanine and RNA guanine bases.*

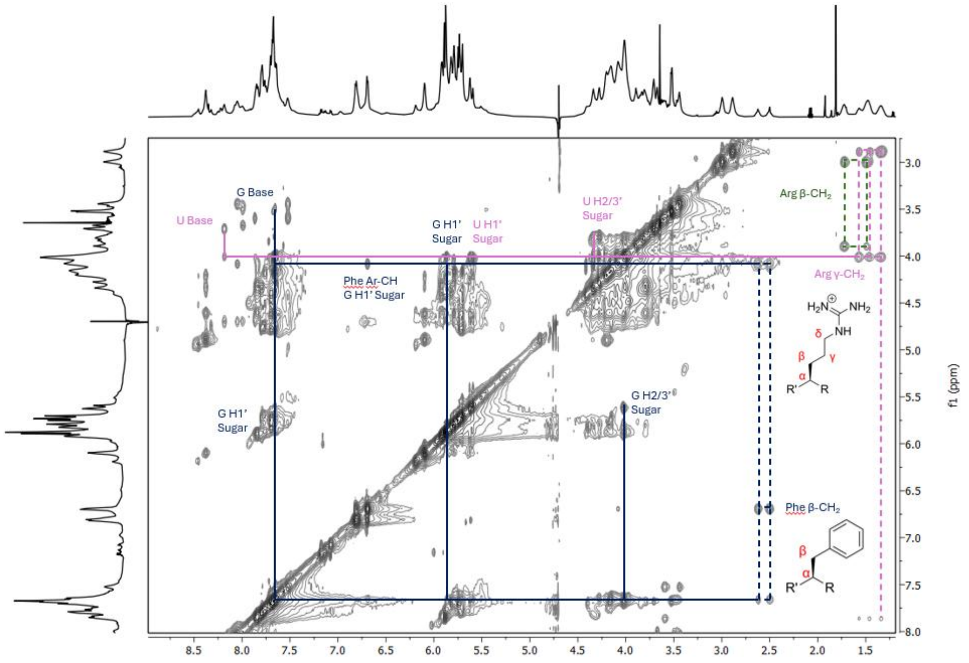

***Supplementary Figure 15:*** *[1]H-[1]H NOESY NMR spectra (600 MHz, 25 °C, 90% $H_2O$, 10% $D_2O$) of* ***RGGFGGRGG*** *peptide and 5'-GGG UUA GGG UUA GGG UUA-3' (TERRA) RNA at pH 7. 2 mM peptide, 2 mM RNA 50 mM Sodium Phosphate (pH 7.01). Interactions: Arginine and RNA Uracil/Guanine Sugars. Phenylalanine and RNA Guanine Sugars.*

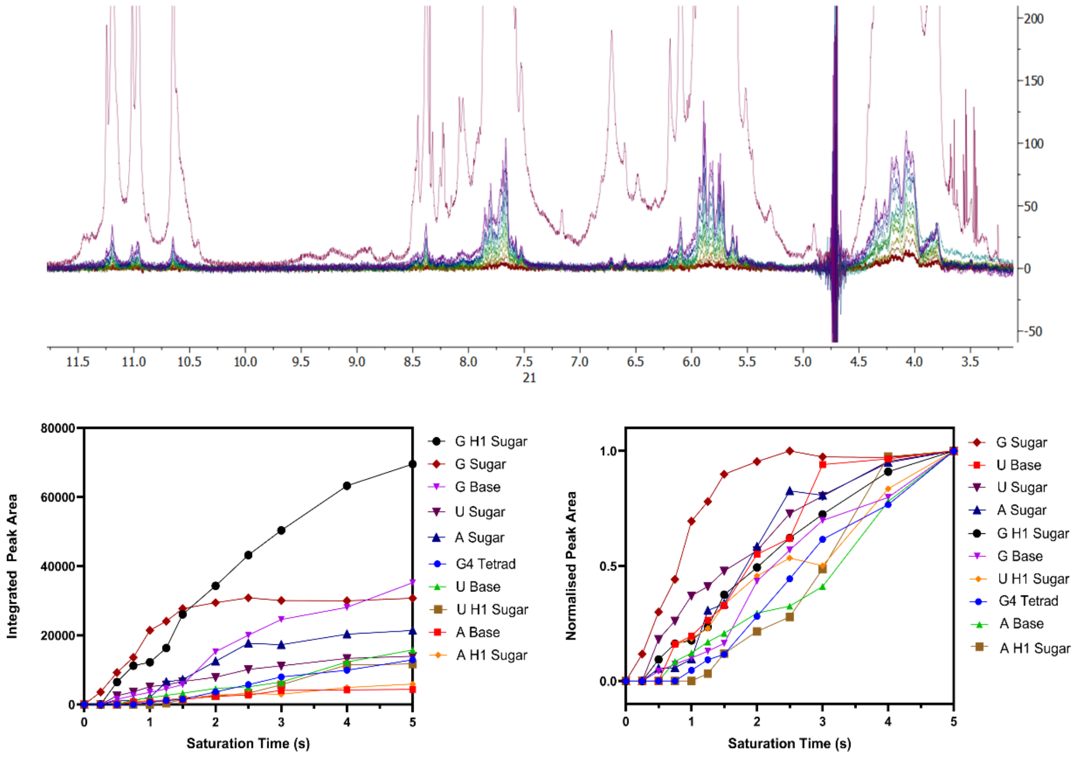

***Supplementary Figure 16:*** *(Top): Saturation Transfer Difference (STD)-NMR spectra of TERRA G-quadruplex (1 mM) &* ***RGGFGGRGG*** *peptide (50 µM). Interactions: Guanine sugars, guanine bases, uracil sugars. 90%* $H_2O$*, 10%* $D_2O$*. 600 MHz. 25 °C. 50 mM Potassium Phosphate (pH 7.01). (Bottom): Left: Raw peak integrations for STD-NMR data. Right: Normalized peak integrations for STD-NMR data.*

#### REFERENCES

1. http://supramolecular.org
2. Brynn Hibbert, D. & Thordarson, P. The death of the Job plot, transparency, open science and online tools, uncertainty estimation methods and other developments in supramolecular chemistry data analysis. *Chemical Communications* **52**, 12792-12805 (2016).
3. Thordarson, P. In *Supramolecular Chemistry: From Molecules to Nanomaterials* (ed P. Gale, Steed, J.), 239-274 (John Wiley & Sons, 2012).
4. von Krbek, L. K. S., Schalley, C. A. &Thordarson, P. Assessing cooperativity in supramolecular systems, *Chem. Soc. Rev.* **46**, 2622-2637 (2017).
5. Schwarz, G. Estimating the Dimension of a Model. *Annals Stat.* **6**, 461-464 (1978).
